# A termination-independent role of Rat1 in cotranscriptional splicing

**DOI:** 10.1101/2020.07.23.218495

**Authors:** Zuzer Dhoondia, Hesham Elewa, Marva Malik, Zahidur Arif, Roger Pique-Regi, Athar Ansari

## Abstract

The yeast termination factor Rat1, and its human homolog Xrn2, have been implicated in multiple nuclear processes. Here we report a novel role of Rat1 in mRNA splicing. Rat1 mutants display increased levels of unspliced transcripts. Accumulation of unspliced transcripts was not due to a failure to degrade unspliced mRNA, disruption of termination or an increased elongation rate in Rat1 mutants. ChIP-Seq analysis revealed Rat1 crosslinking to the introns of a subset of yeast genes. Mass spectrometry and coimmunoprecipitation revealed interaction of Rat1 with the Clf1, Isy1, Yju2, Prp43, and Sub2 splicing factors. Furthermore, recruitment of the Prp2 splicing factor on the intron was compromised in the Rat1 mutant. Based on these findings we propose that Rat1 has a novel role in splicing of a subset of mRNA in budding yeast.

## Introduction

Rat1/Xrn2 are 5′→3′ exoribonucleases belonging to the Xrn-family of nucleases (Nagarajan et al., 2013). These are highly conserved proteins with homologs present in budding yeast, fission yeast, flies, plants, worms, mice and humans (Johnson, 1997; Kastenmayer and Green, 2000; Richter et al., 2016; Shobuike et al., 1995; Sugano et al., 1994; Zhang et al., 1999). Rat1/Xrn2 have been implicated in multiple aspects of RNA metabolism in eukaryotes. Studies carried out in different eukaryotic systems have demonstrated involvement of Rat1/Xrn2 in RNA trafficking, RNA quality control, RNA processing, promoter-associated transcription, elongation and termination of transcription (Amberg et al., 1992; Bousquet-Antonelli et al., 2000; Brannan et al., 2012; Jiao et al., 2010; Jimeno-González et al., 2010; Kim et al., 2004; Miki et al., 2017; West et al., 2004).

Rat1 was discovered in budding yeast as a factor required for nucleo-cytoplasmic transport of mRNA. In a temperature-sensitive mutant of Rat1, mRNA was cleaved and polyadenylated but failed to move out of the nucleus and hence was named <u>R</u>N<u>A</u> <u>t</u>rafficking factor or Rat1 (Amberg et al., 1992). Further investigation revealed that Rat1 and its higher eukaryotic homolog Xrn2 are involved in termination of RNAPII-mediated transcription in a manner dependent on their 5′→3′ exoribonucleases activity (Kim et al., 2004; West et al., 2004). The termination of transcription involves pausing of RNAPII beyond the poly(A)-site, which facilitates recruitment of CF1 and CPF 3’ end processing/termination factors in yeast (Mischo and Proudfoot, 2013). The transcribing mRNA is cleaved downstream of the poly(A)-site by the Ysh1 subunit of CPF complex and is polyadenylated by poly(A)-polymerase (Butler and Platt, 1988; Jenny et al., 1996). The monophosphorylated mRNA downstream of the cleavage site is still attached to the elongating RNAPII. Rat1 bind to the uncapped transcript and the 5′→3′ exoribonucleases activity digests the nascent RNA. Removal of the RNA helps disengage RNAPII from the DNA template (Kim et al., 2004). The Rat1 homolog Xrn2 facilitates termination of transcription in higher eukaryotes by a similar mechanism (Eaton et al., 2018; Fong et al., 2015; West et al., 2004). Rat1/Xrn2-mediated termination is thus reminiscent of the ‘torpedo’ mechanism of termination of transcription in prokaryotes (Jin et al., 1992). The enzymatic activity of Rat1 is crucial but not sufficient for termination of transcription (Pearson and Moore, 2013). Genomewide studies have revealed that Rat1 is not a universal termination factor like CF1 subunit Pcf11 and CPF subunit Ysh1, but is required for termination of transcription of only 35% of RNAPII-transcribed genes in budding yeast (Baejen et al., 2017).

Rat1 may also be involved in termination at the pre-cleavage/polyadenylation step in accordance with the ‘allosteric’ mechanism. The pausing of the polymerase near the poly(A)-site, which is critical for both cleavage/polyadenylation and termination, is dependent on Rat1. In the *rat1-1* mutant, which exhibits a termination defect, there is hyperphosphorylation of CTD-serine-2 resulting in an increased elongation rate. The *rat1-1* termination defect could be rescued by introduction of the *rpb1-N488D* allele, which causes a slower elongation rate (Jimeno-González et al., 2010). These findings strongly suggest that Rat1 has dual roles in termination; a pre-cleavage role in RNAPII pausing downstream of the poly(A)-site, and a post-cleavage role in degrading 5′ monophosphorylated RNA followed by dissociation of RNAPII from the DNA template. Another important implication is involvement of Rat1 in the elongation step of transcription through its influence on hyperphosphorylation of the CTD at serine-2 (Jimeno-González et al., 2014). Indeed, combining *rat1-1* allele with the *rpb1-E1103G* mutation, which causes a faster polymerase elongation rate, resulted in an enhanced growth defect (Jimeno-González et al., 2010; Malagon et al., 2006). Depletion of Xrn2 in human cell lines resulted in promoter-proximal pausing, which also could be the consequence of the role of the protein in early elongation step (Nojima et al., 2015).

In multiple organism Rat1 function appears linked to the promoter-terminator crosstalk. In *C. elegans*, not all genes that exhibit 3′ end occupancy of Rat1 are dependent on this protein for termination. Whether Rat1 is required for termination is determined by the promoter element in worms (Miki et al., 2017). In budding yeast also, Rat1 could dismantle only the promoter-driven transcription complexes in an *in vitro* assay (Pearson and Moore, 2013). In keeping with a role in gene looping, Rat1/Xrn2 have been found to bind both the 5′ and 3′ ends of genes (Baejen et al., 2017; Fong et al., 2015; Kim et al., 2004; Nojima et al., 2015). Localization of Rat1/Xrn2 in the promoter-proximal region gave rise to speculation that it plays a role in promoter-associated transcription. Experimental evidence has implicated Rat1/Xrn2 in the initiation-elongation transition (Harlen and Churchman, 2017). In the absence of Xrn2, a peak of promoter-proximally paused polymerase was observed, thereby suggesting that Xrn2 is involved in early stages of transcription. Xrn2 has also been found to function in promoter directionality as an increase in polymerase signal was observed in the region upstream of promoters in the absence of Xrn2 activity (Brannan et al., 2012; Fong et al., 2015). A similar involvement of Rat1 in promoter directionality in budding yeast cannot be ruled out. Both Rat1 and Xrn2 have also been shown to play a role in mRNA quality control by degrading aberrant uncapped transcripts (Brannan et al., 2012; Jiao et al., 2010; Jimeno-González et al., 2010). In line with these results, human Xrn2 has been reported to coimmunoprecipitate with the Edc3, Dcp1a and Dcp2 capping proteins (Brannan et al., 2012). No such interaction of Rat1 with capping enzymes has been observed in budding yeast, but Rat1 can similarly degrade uncapped yeast transcripts (Jiao et al., 2010).

Rat1/Xrn2 have also been implicated in degradation of splicing-defective transcripts. Xrn2 was reported to degrade unspliced transcripts in human cells, while Rat1 in yeast was linked to degradation of unspliced transcripts in the splicing factor mutant Prp2 (Bousquet-Antonelli et al., 2000). We discovered accumulation of unspliced transcripts in mutants of Rat1 with a wild type Prp2 allele. This increase was not due to stabilization of unspliced transcripts, or an indirect effect of defective termination, or due to an absence of polymerase pausing on intronic regions in the Rat1 mutant. We present evidence that Rat1 plays a direct role in splicing of precursor mRNA. We discovered physical interaction of Rat1 with the intronic sequences as well as with the splicing factors of NineTeen complex (NTC). Furthermore, recruitment of the Prp2 splicing protein to the intron was compromised in the absence of functional Rat1. Our findings strongly suggest a novel role of Rat1 in splicing of primary transcripts of a subset of genes in budding yeast.

## Results

### Unspliced mRNA accumulates in the absence of functional Rat1

Rat1 has been implicated in a variety of nuclear processes in yeast and higher eukaryotes. In order to have a comprehensive understanding of the role of Rat1 in RNA biogenesis in budding yeast, we monitored mRNA levels of selected genes in the temperature-sensitive *rat1-1* mutant. RNA analysis was performed in cells grown at the permissive (25°C) and non-permissive (37°C) temperatures by RT-PCR. Briefly, the protocol involved isolation of total RNA from exponentially growing cells, reverse transcription of RNA using an oligo-dT, PCR amplification of resultant cDNA using gene-specific primers and quantification of RT-PCR products as described in El Kaderi et al., (2009). Since Rat1 affects transcription of several mRNA species as well as 5.8S and 28S rRNAs, we used 5S rRNA as normalization control in all experiments. In accord with the termination function of Rat1, a decrease in mRNA level of most genes was observed upon shifting of cells to the non-permissive temperature (Figure 1B). However, we observed an unusual result with the *APE2* gene. A longer transcript appeared upon shifting of *rat1-1* cells to the non-permissive temperature (Figure 1B, lane 4). *APE2* is one of the few intron-containing genes in budding yeast, and sequencing of the PCR product revealed that the longer transcript was the unspliced mRNA. To determine if the accumulation of unspliced transcripts is unique to *APE2*, we examined several other intronic genes. Of all the genes that we analyzed, *STO1*, *ASC1* and *IMD4,* exhibited a similar trend of increased unspliced mRNA in the *rat1-1* mutant at the elevated temperature (Figure 1B, lane 4 and Figure 1C, black bars). No such accumulation of unspliced transcripts was observed in the isogenic wild type cells at 37°C (Figure 1B, lane 2 and Figure 1C). RT-PCR performed using gene-specific primers gave identical results. Not all intron-containing genes, however, exhibit increased unspliced transcript level in the mutant. Out of eight genes that we tested, four exhibited splicing defects.

**Figure 1.**
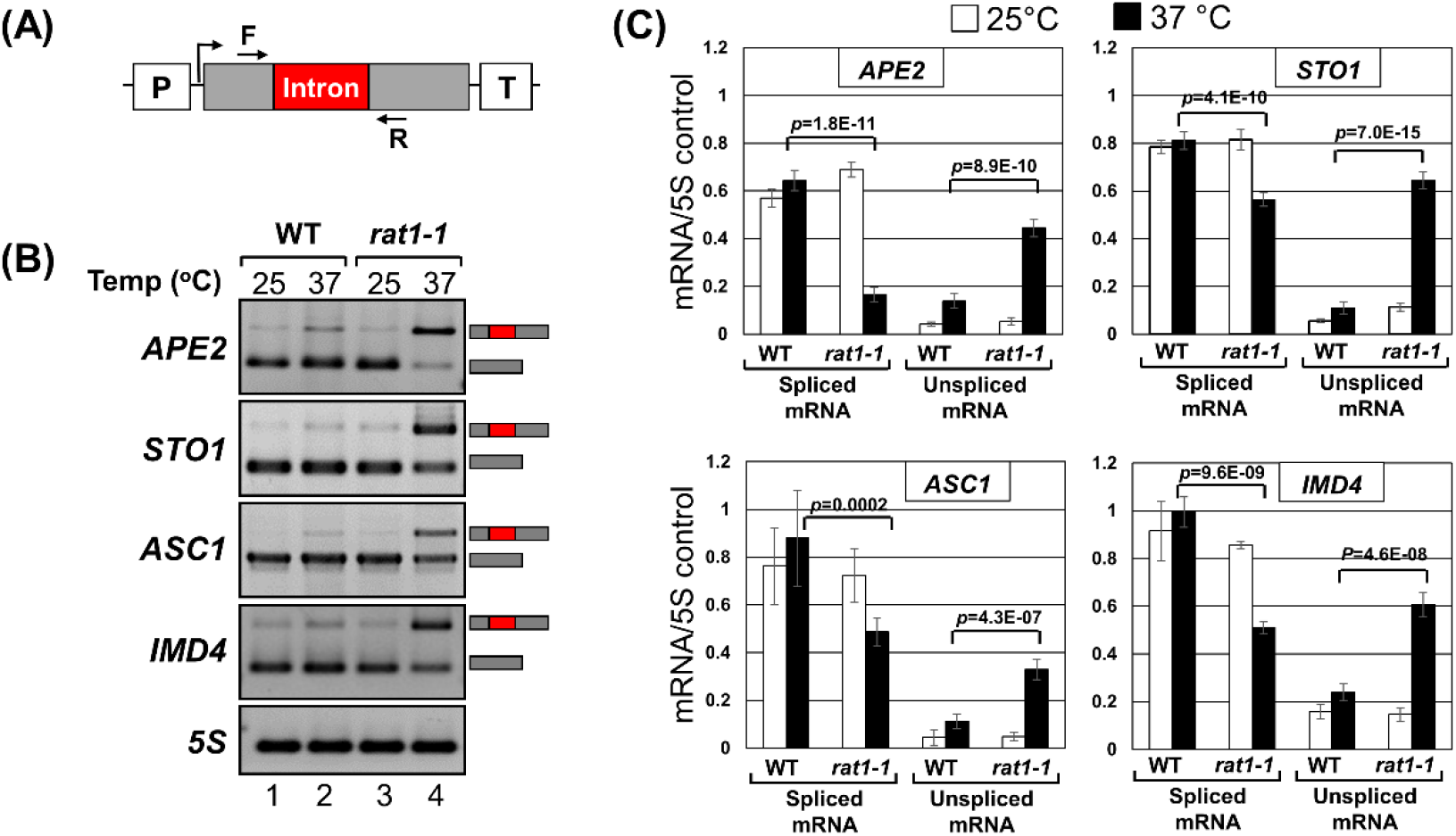
Unspliced transcripts accumulate in the *rat1-1* mutant at non-permissive temperature. (A) Schematic depiction showing the position of primers F and R used in RT-PCR analysis. (B) RT-PCR analysis of mRNA in wild type (WT) and *rat1-1* mutant cells at the indicated temperatures. (C) Quantification of data shown in (B). 5S rRNA was used as normalization control. The error bars represent one unit of standard deviation. The results shown are an average of six independent PCRs from three separate RNA preparations from three independently grown cultures.

To confirm that accumulation of unspliced transcripts in the *rat1-1* strain was due to the inactivation of Rat1, and not due to a factor in the genetic background of the *rat1-1* strain, we used the ‘anchor away’ approach. This technique involves selective depletion of a protein from the nucleus by anchoring it to ribosomes in the cytoplasm (Haruki et al., 2008). The approach takes advantage of the rapamycin-dependent heterodimerization of FK506 binding protein (FKBP12) with FKBP12-rapamycin-binding (FRB) domain (Belshaw et al., 1996; Chen et al., 1995). We inserted the FRB domain at the carboxy-terminus of Rat1 in a strain that has FKBP12 fused to the ribosomal protein RPL13A, a *fpr1* deletion and a *tor1-1* mutation. The resultant strain was named *Rat1-AA*. In the presence of rapamycin, Rat1-FRB dimerized with ribosomal RPL13A-FKBP12 in the anchor away strain, leading to almost complete depletion of Rat1 from the nucleus within 60 minutes of addition of the antibiotic to the medium. We monitored levels of spliced and unspliced transcripts in the *Rat1-AA* strain in the presence of rapamycin (Rat1 depleted from the nucleus) and in the absence of rapamycin (Rat1 present in the nucleus) by RT-PCR as described above. We observed a 2-4-fold increase in the level of unspliced *APE2, STO1, MRK1, NMD2* and *LSB4* transcripts upon depletion of Rat1 (Figure 2C, lane 4 and Figure 2D, red bars). There was no such rapamycin-dependent buildup of unspliced mRNA in the isogenic wild type strain (Figure 2C, lane 2 and Figure 2D, red bars). We conclude that there is an accumulation of unspliced transcripts when Rat1 is inactivated or depleted from the nucleus. All intron-containing genes of budding yeast, however, do not exhibit accrual of unspliced RNA in the absence of Rat1. Out of eight genes that we tested, only four exhibited accumulation of unspliced transcripts in the absence of Rat1, while four showed no difference.

**Figure 2.**
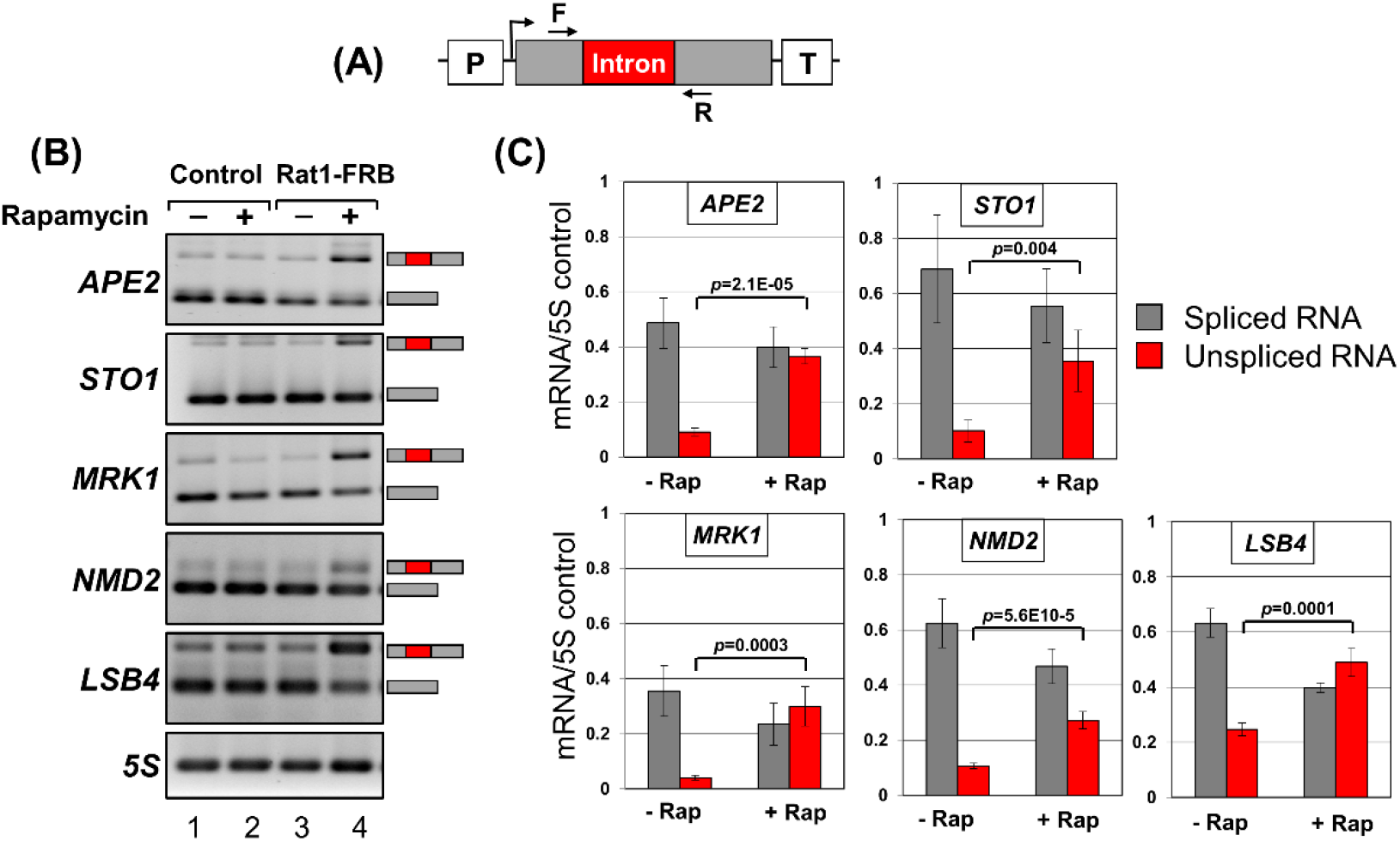
Accumulation of unspliced transcripts upon nuclear depletion of FRB-tagged Rat1 in the presence of rapamycin. (A) Schematic depiction of a gene showing the position of primers F and R used in RT-PCR analysis. (B) Gel pictures showing RT-PCR products for the indicated genes in FRB-tagged Rat1 strain and untagged strain (control) in the presence and absence of rapamycin. (C) Quantification of data in lanes 3 and 4 of (B). 5S rRNA was used as normalization control. The error bars represent one unit of standard deviation. The results shown are an average of six independent PCRs from three separate RNA preparations from three independently grown cultures.

To explain the presence of unspliced mRNA in the absence of Rat1 activity, we formulated four hypotheses: (1) Rat1 plays a surveillance role and degrades unspliced transcripts by its exoribonuclease activity; (2) accumulation of unspliced transcripts is the consequence of an indirect effect of Rat1-mediated termination on splicing; (3) Rat1 effect on elongation facilitates pausing of the polymerase during transcription of intronic regions to enable completion of splicing reactions; (4) Rat1 has a novel role in splicing of primary transcripts.

We tested each of these hypotheses and the results are described below.

### Rat1 is not the surveillance factor that degrades unspliced transcripts

The experiments described above monitored steady-state RNA levels, which is the net product of two opposing processes, transcription and degradation. To determine if Rat1 has a surveillance role in degrading unspliced mRNA by its exoribonuclease activity, we measured levels of unspliced and spliced transcripts by the transcription run-on (TRO) approach. The TRO assay detects nascent transcripts and therefore reflects transcription and rules out the contribution of RNA stability to the measured RNA content. The strand-specific TRO analysis was carried out using Br-UTP by the modification of the method described in Core et al., (2008). Briefly, exponentially growing yeast cells were permeabilized with sarkosyl and allowed to resume transcription in the presence of Br-labelled UTP for 2-5 minutes. Br-UTP-labelled nascent RNA was immunopurified, reverse transcribed using oligo-dT and amplified by PCR as described in Dhoondia et.al (2017). TRO analysis revealed a 2-7-fold increase in the levels of unspliced transcripts of *APE2*, *STO1*, *MRK1* and *LSB4* upon depletion of Rat1 in the *Rat1-AA* strain (Figure 3B and C).

**Figure 3.**
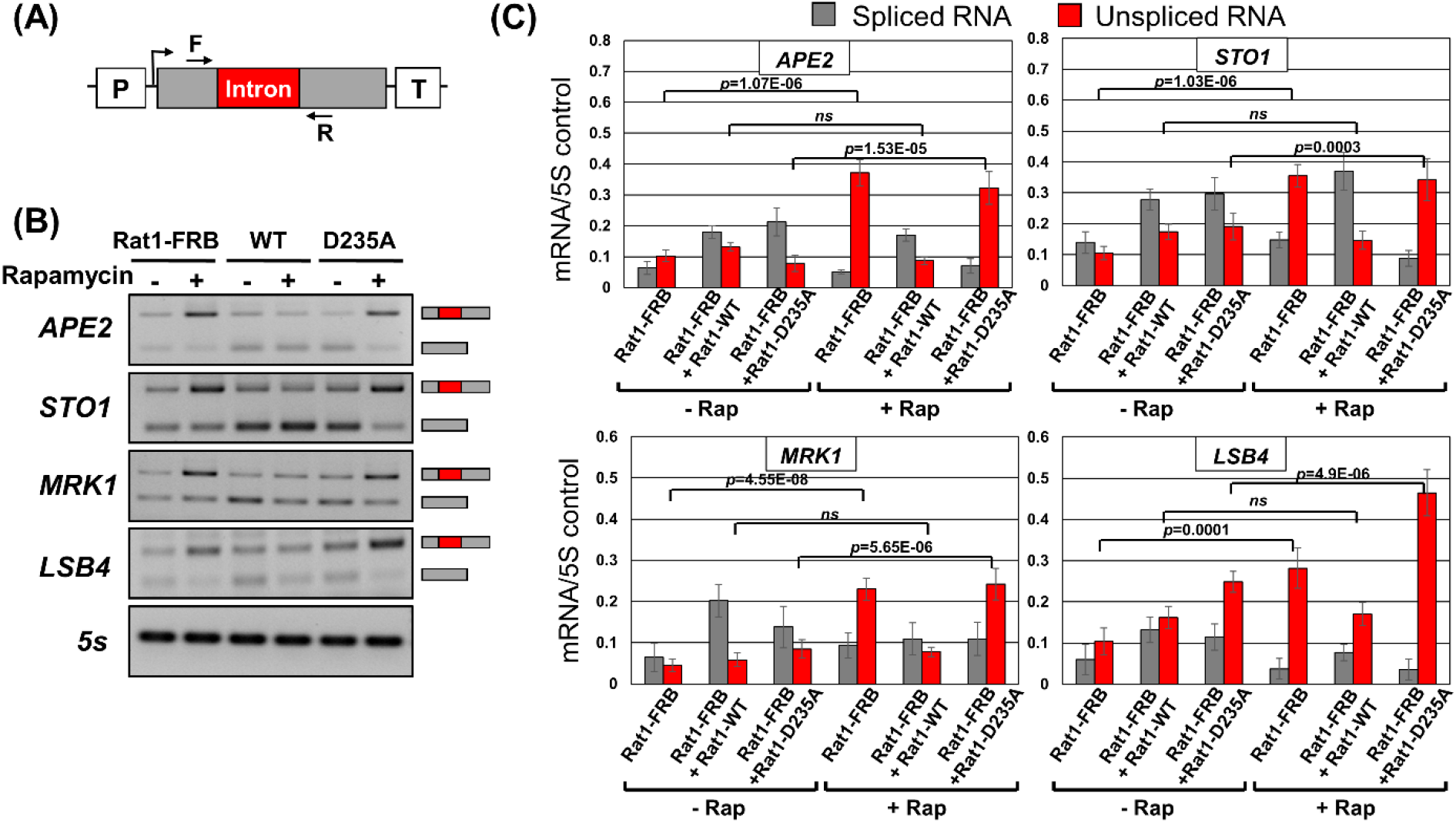
Transcription Run-On (TRO) analysis showing that accumulation of nascent unspliced transcripts upon nuclear depletion of FRB-tagged Rat1 in the presence of rapamycin is rescued by wild type Rat1 but not the catalytically inactive mutant. (A) Schematic depiction of a gene showing the position of primers F and R used in TRO analysis. (B) Quantification of TRO signals for the indicated genes in FRB-tagged Rat1 strain in the presence and absence of rapamycin and upon complementation with WT Rat1 or catalytically inactive D235A Rat1 mutant. 5S rRNA was used as normalization control. The error bars represent one unit of standard deviation. The results shown are an average of six independent PCRs from three separate RNA preparations from three independently grown cultures.

Since splicing occurs cotranscriptionally in both yeast and higher eukaryotes, these findings suggest that Rat1 may be affecting cotranscriptional splicing of mRNA. The possibility of Rat1 playing a role in cotranscriptional degradation of unspliced transcripts, however, could not be ruled out. To resolve the issue, we transformed the *Rat1-AA* strain with a plasmid expressing either wild type Rat1 or a mutant form of Rat1 containing a point mutation at an evolutionarily conserved catalytic residue (D235A). Denoted as exo-Rat1, the mutant is deficient in exoribonuclease activity (Kim et al., 2004). TRO was repeated in the presence and absence of rapamycin. Expression of the wild type Rat1 rescued the accumulation of unspliced transcript, but the catalytically inactive exo-Rat1 mutant was unable to do so (Figure 3B and 3C). One interpretation is that Rat1 catalytic activity is required for degradation of unspliced transcripts, and therefore there is a buildup of unspliced mRNA in the exo-Rat1 mutant.

Rat1 is highly specific for 5′ monophosphorylated RNA substrates. 7′-methylguanosine capped mRNA is not a suitable substrate of the Rat1 nuclease. Thus, Rat1 will be able to degrade unspliced mRNA only if it is uncapped. We therefore examined the capping pattern of unspliced transcripts. RNA was isolated from mutant *rat1-1* cells grown at 25°C and 37°C and subjected to immunoprecipitation (RNA-IP) using antibodies against the 7′-methylguanosine cap. Immunoprecipitated RNA was subjected to RT-PCR for *APE2*, *STO1*, *RPS14A* and *RPS9B* as described previously. 5S rRNA was used as a normalization control to ensure that equal amounts of RNA were used for immunoprecipitation. As expected, there was an approximate 7-fold increase in the amount of unspliced mRNA of *APE2*, *STO1*, *RPS14A* and *RPS9B* at 37°C compared to 25°C (Figure 4B) and there was no accumulation of unspliced transcripts with 7′-methyl guanosine caps in the wild type strain (Figure 4–figure supplement 1). About 70 to 80% of both spliced and unspliced transcripts of the four genes analyzed could be immunoprecipitated by anti-7′-methylguanosine antibodies (Figure 4B). This demonstrates that the degree of capping of unspliced mRNA is similar to that of spliced mRNA, and therefore Rat1 enzymatic activity is unlikely to be directed to unspliced transcripts by decapping. We conclude that the accumulation of unspliced mRNA in the absence of Rat1 activity is not likely due to a surveillance role of Rat1 in degrading intron-containing transcripts.

**Figure 4.**
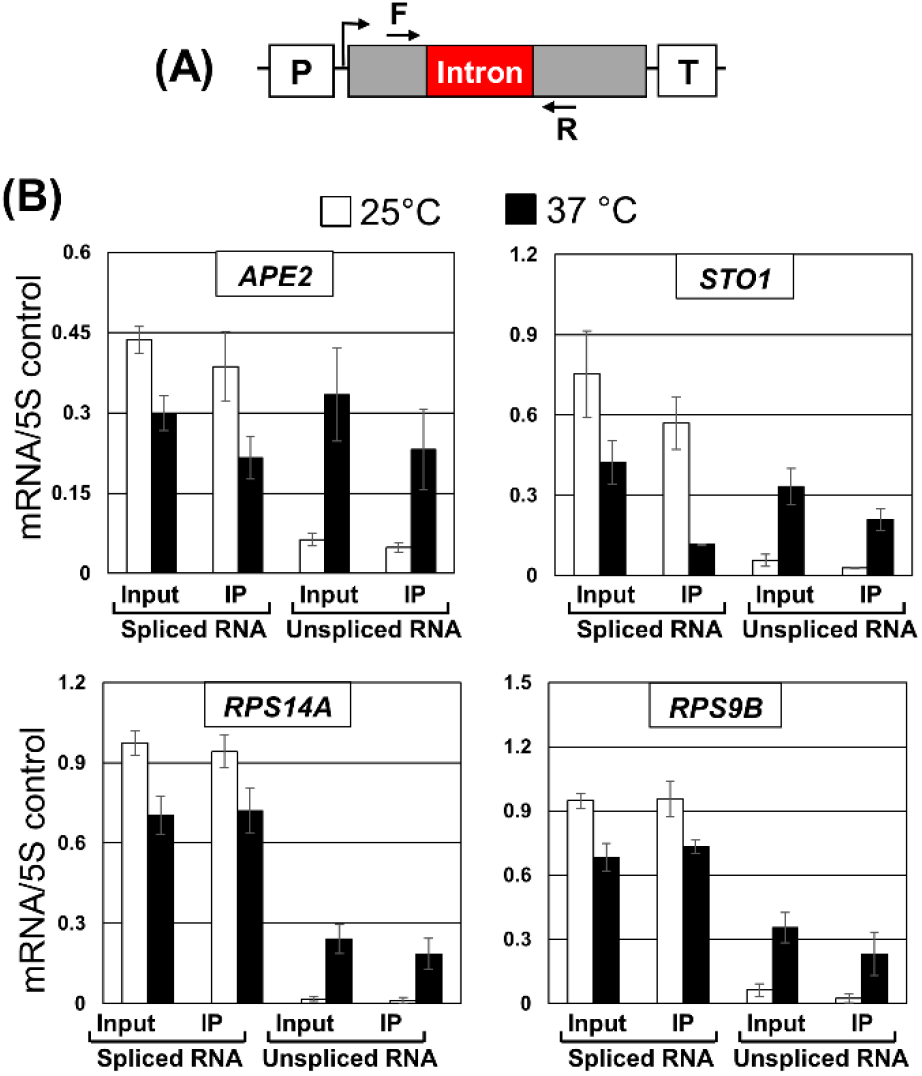
RNA immunoprecipitation analysis shows that unspliced transcripts in *rat1-1* mutant are capped. (A) Schematic depiction of a gene showing the position of primers F and R used in RT-PCR analysis. (B) Affinity purified RNA using anti-m7G were reverse transcribed. Quantification of data shown in for the indicated genes in *rat1-1* mutant at the indicated temperatures. 5S rRNA was used as normalization control. The error bars represent one unit of standard deviation. The results shown are an average of six independent immunoprecipitates from three separate RNA preparations from three independently grown cultures.

### Accumulation of unspliced mRNA in the absence of Rat1 is not the consequence of defective termination

Rat1 and its homolog Xrn2 are transcription termination factors. We reasoned that if appearance of unspliced transcripts in the absence of Rat1 activity is due to an indirect effect of defective termination on splicing, other termination-defective mutants may also display a splicing defect.

Termination of transcription by RNAPII in yeast requires the CF1 and CPF complexes. Both complexes are required for cleavage and polyadenylation of the messenger. The cleavage-polyadenylation of primary transcripts is coupled to the poly(A)-dependent termination of transcription. A recent genome wide analysis revealed that Rat1 is required for termination of nearly 35% of yeast genes (Baejen et al., 2017). In contrast, CF1 and CPF subunits have a more robust role as they affect poly(A)-dependent termination of nearly 90% of genes (Baejen et al., 2017). To determine if accumulation of unspliced transcripts is an indirect effect of defective termination in the Rat1 mutant, we monitored levels of spliced and unspliced mRNA of *APE2*, *STO1*, *ASC1* and *IMD4* in the termination-defective *rna14-1* mutant. Rna14 is a subunit of CF1 complex and is essential for poly(A)-dependent termination (Minvielle-Sebastia et al., 1997). There was a decrease in the level of polyadenylated mRNA of *APE2*, *STO1*, *ASC1* and *IMD4* upon shifting of mutant cells to the elevated temperature (Figure 5B, lane 4 and Figure 5C, black bars) in agreement with the need for termination to achieve optimal transcription. There was, however, no increase in the level of unspliced transcripts at the non-permissive temperature (Figure 5B, lane 4 and Figure 5C, black bars). The amount of unspliced mRNA in the mutant at 37°C was similar to that in the isogenic wild type strain (Figure 5B, lanes 2 and 4 and Figure 5C, black bars). The defective termination in the Rna14 mutant therefore did not have an adverse effect on splicing. The accumulation of unspliced mRNA in the Rat1 mutant is thus unlikely to be the consequence of defective termination.

**Figure 5.**
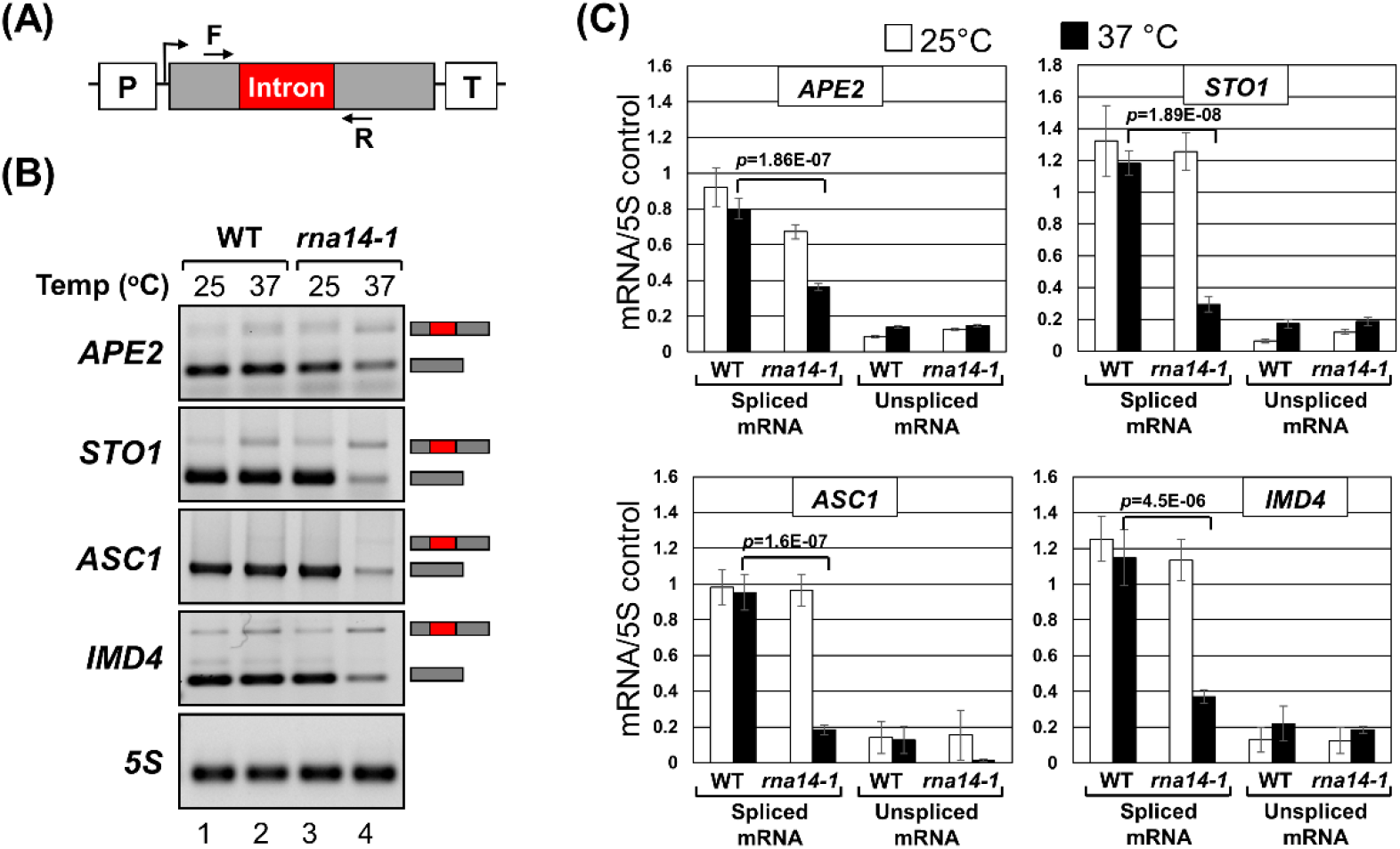
Defective termination in *rna14-1* mutant does not result in accumulation of unspliced transcripts. (A) Schematic depiction of a gene showing the position of primers F and R used in RT-PCR analysis. (B) Gel pictures showing RT-PCR products for the indicated genes in wild type (WT) and *rna14-1* mutant at the indicated temperatures. (C) Quantification of data shown in (B). 5S rRNA was used as normalization control. The error bars represent one unit of standard deviation. The results shown are an average of at least six independent PCRs from three separate RNA preparations from three independently grown cultures.

### Does Rat1 facilitated pausing of polymerase during elongation of intronic regions enable the recruitment of splicing factors?

During cotranscriptional splicing, the polymerase slows down or pauses while transcribing intronic regions (Braberg et al., 2013). This may enable recruitment of splicing factors, leading to efficient execution of the splicing reaction. Consequently, the increased elongation rate associated with some mutants of RNAPII produces splicing defects (Braberg et al., 2013). The *rat1-1* mutant also exhibits enhanced RNAPII elongation rate by stimulating hyper-phosphorylation of CTD-serine-2 (Jimeno-González et al., 2014). The possibility of elongation speed contributing to the splicing defect observed in *rat1-1* mutant thus could not be ruled out.

We therefore examined pausing of polymerase in the absence of Rat1 activity using Rpb1-ChIP-Seq data from Baejen et al., (2017). This study analyzed genome wide transcription using Rpb1-ChIP-Seq approach following Rat1 depletion from the nucleus. We extracted Rpb1-ChIP-Seq data for 280 intron containing genes analyzed in this study. Rpb1-ChIP-Seq reads were aligned with respect to 5′ and 3′ splice sites of introns and included a 400 bp window upstream of the 5′ splice site and a 1 kbp window downstream of the 3′ splice site (Figure 6). There was no global change in polymerase density over the intron upon nuclear depletion of Rat1 (Figure 6A and Figure 6B). Rpb1-ChIP profiles of individual genes confirms that Rpb1 signal over the intron and splice sites is similar (Figure 6C). As expected, there was an increased RNAPII read-through signal in 1 kbp window downstream of termination sites upon nuclear depletion of Rat1, confirming a termination defect. These findings indicate that accumulation of unspliced transcripts in Rat1 mutants is not an indirect consequence of loss of pausing of the polymerase over intronic regions in the Rat1 mutant.

**Figure 6.**
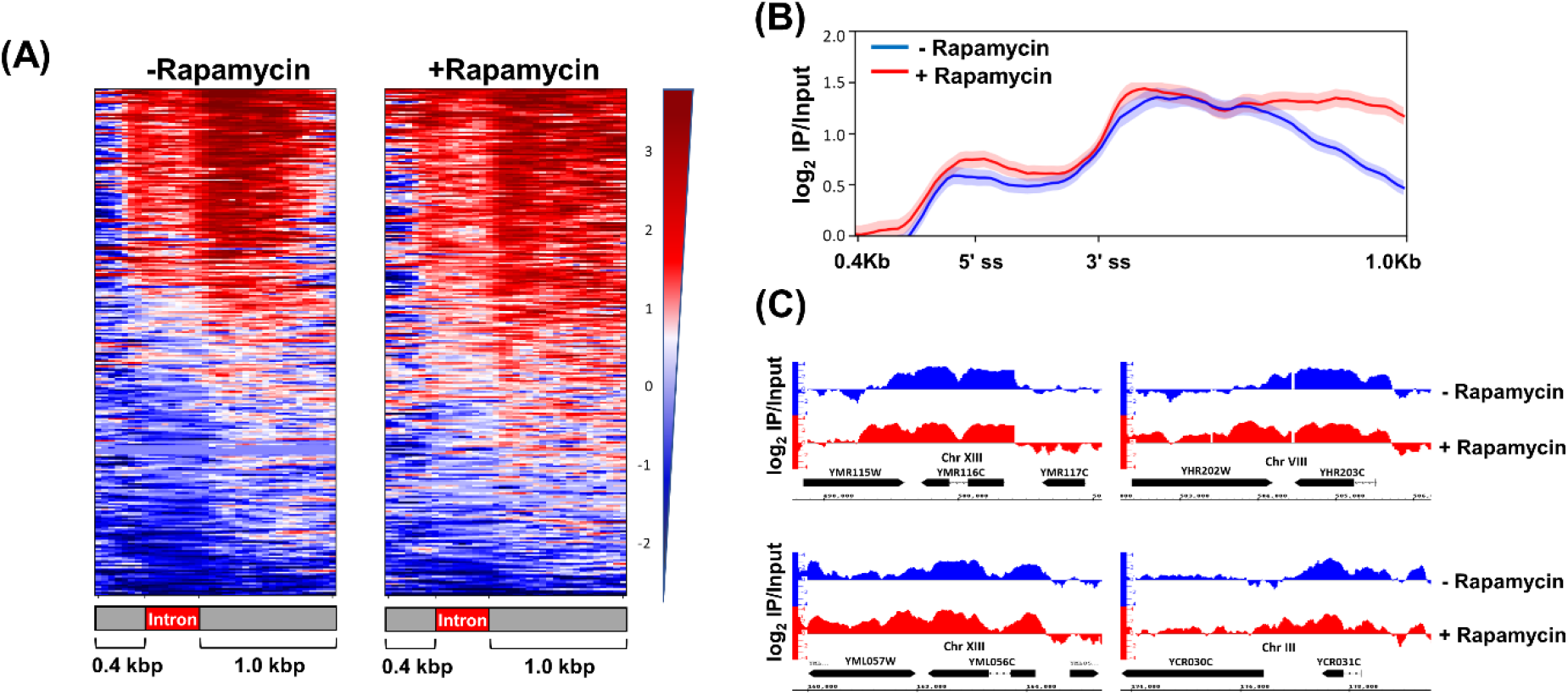
RNAPII occupancy across the intronic genes after nuclear depletion of Rat1. (A) Heatmaps of Rpb1 occupancy levels calculated as the log2 IP/Input ratio in Rat1-FRB strain before (−rapamycin) and after (+rapamycin) nuclear depletion of Rat1 from ChIP-seq experiments performed in Baejen et.al., 2017. Genes are sorted in the descending order of Rpb1 occupancy. The y-axis indicates individual genes from the 280 intron containing genes and the x-axis shows relative position of 0.4 Kbp upstream and 1.0 Kbp downstream around 5′ ss and 3′ ss intronic sites. (B) Averaged normalized occupancy profiles from the ChIP-seq data of RNAPII. Metagene plot was designed with intron size averaged to 300 bp and plotting 0.4 Kbp upstream and 1Kbp downstream. The profiles are aligned at both the 5′ ss and 3’ ss of 280 intron containing genes. The shaded areas around the ChIP-seq profile represent error ±1. (C) IGB browser view of log2 IP/Input Rpb1 (RNAPII) read counts from ChIP-seq experiment performed in Rat1-FRB tagged strain before (blue) and after (red) nuclear depletion of Rat1. Four representative ORFs shown are *YMR116C, YHR203C, YML056C, YCR031C*.

### Rat1 has a novel role in splicing of mRNA

None of the studies described above provide an explanation for accumulation of unspliced transcripts in the absence of functional Rat1. An alternative possibility is that Rat1 has a novel role in splicing of primary transcripts. Therefore, we examined the direct involvement of Rat1 in splicing by three parallel approaches. We reasoned that if Rat1 is involved in splicing, (1) it will physically interact with introns; (2) it will exhibit a transient or stable interaction with the splicing factors; and (3) it will facilitate the recruitment of splicing factors on the intron.

### Rat1 is enriched over introns

We performed ChIP to determine if Rat1 is enriched over intronic DNA. A TAP-tag was inserted at the carboxy-terminus of Rat1. TAP-tag does not interfere with the termination function of Rat1. We used this strain for high resolution ChIP that employs extensive sonication as described in El Kaderi et al., (2009). Splicing factors do not directly crosslink to the splice sites on DNA but interactions with the elongating transcript and CTD of RNAPII during cotranscriptional splicing provides sufficient indirect contacts to detect splicing factors by ChIP on DNA (Herzel et al., 2017; Nojima et al., 2018). To favor indirect contacts we used a more robust crosslinking approach, disuccinimidyl glutarate (DSG) together with formaldehyde, as described in GRID-Seq approach (Li et al., 2017). We found that Rat1 crosslinks to the promoter and terminator regions of *APE2* (Figure 7–figure supplement 1, primers pairs A and E) in accordance with published findings (Baejen et al., 2017; Kim et al., 2004). More importantly, Rat1 also localized to the intronic splice sites of *APE2* (Figure 7–figure supplement 1, primer pair C). The crosslinking of Rat1 to the splice sites suggest that Rat1 may have a direct role in splicing of *APE2* mRNA.

We next examined if other intron-containing genes similarly exhibit crosslinking of Rat1 to the intron. We used ChIP-Seq data from Baejen et al., (2017) for this analysis. We extracted data for intron-containing genes from this analysis. The Rat1 occupancy profile on 280 intron-containing genes, aligned to TSS and TES ± 400 bp, showed that Rat1 is localized at the terminator and promoter regions of genes (Figure 7–figure supplement 2A). We also analyzed occupancy of Pcf11, which is an essential termination factor and a component of CF1A 3′ end processing/termination complex, at TSS and TES as described above using the data from the same genomewide study. Pcf11 and Rat1 occupancy profile were comparable at the termination region. They, however, differed considerably near the promoter. Rat1 occupancy was significantly higher than that of Pcf11 just downstream of the promoter (Figure 7–figure supplement 2A). Most yeast introns are positioned near the TSS and the specific enrichment of Rat1 near the TSS might be due to its localization to the intron. We then examined the Rat1 and Pcf11 occupancy profiles of non-intronic genes and found them essentially identical (Figure 7–figure supplement 2B). These suggests that the Rat1 enrichment at the TSS may be due in part to Rat1 crosslinking to the promoter-proximal intron.

**Figure 7.**
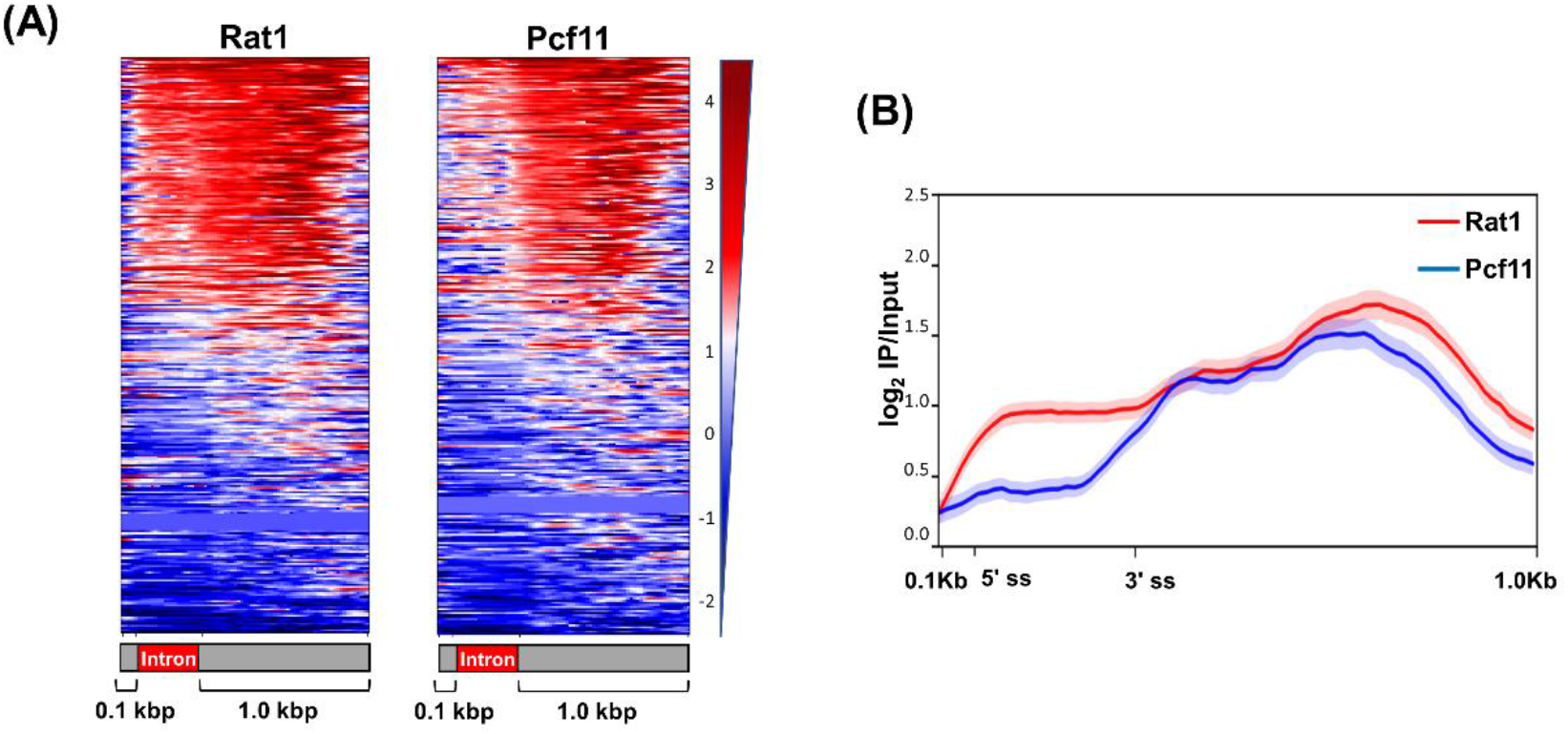
Rat1 is selectively enriched in the intronic region of the genes. (A) Heatmaps of Rat1 and Pcf11 occupancy levels calculated as the log2 IP/Input ratio from ChIP-Seq experiments performed in Rat1-TAP and Pcf11-TAP strains, respectively. Genes are sorted in the descending order of the occupancy. The y-axis indicates individual genes from the 280 intron containing genes and the x-axis shows relative position of 0.1 Kbp upstream and 1.0 Kbp downstream around 5′ ss and 3′ ss intronic sites. (B) Averaged normalized occupancy profiles from the ChIP-Seq data of Rat1 (red) and Pcf11 (blue). The metagene plot was designed with an average intron size of 300 bp and plotting 0.1 Kbp upstream and 1Kbp downstream. The profiles are aligned at both the 5′ ss and 3′ ss of 280 intron containing genes. The shaded areas around the ChIP-Seq profile represent standard error ±1.

Next, Rat1-ChIP-Seq reads were aligned with respect to the 5′ and 3′ splice sites of introns, encompassing a 100 bp window upstream of the 5′ splice sites and a 1 kbp window downstream of the 3′ splice sites as shown in Figure 7. Out of 280 intron-containing genes analyzed, 105 exhibited crosslinking of Rat1 to the intronic sequence. Thus, not all intron-containing genes show Rat1 occupancy. Unlike Rat1, Pcf11 does not exhibit significant localization to the intronic region (Figure 7A and 7B). These results indicate that Rat1 localizes to the introns of a subset of genes in budding yeast. We randomly selected five of the 105 intron-containing genes that exhibited Rat1-intron occupancy to test whether Rat1 occupancy correlated with Rat1-dependence on splicing. Out of these five genes, four were dependent on Rat1 for efficient splicing (Figure 8–figure supplement 1). Thus, not all introns that recruit Rat1 are dependent on Rat1 for splicing.

**Figure 8.**
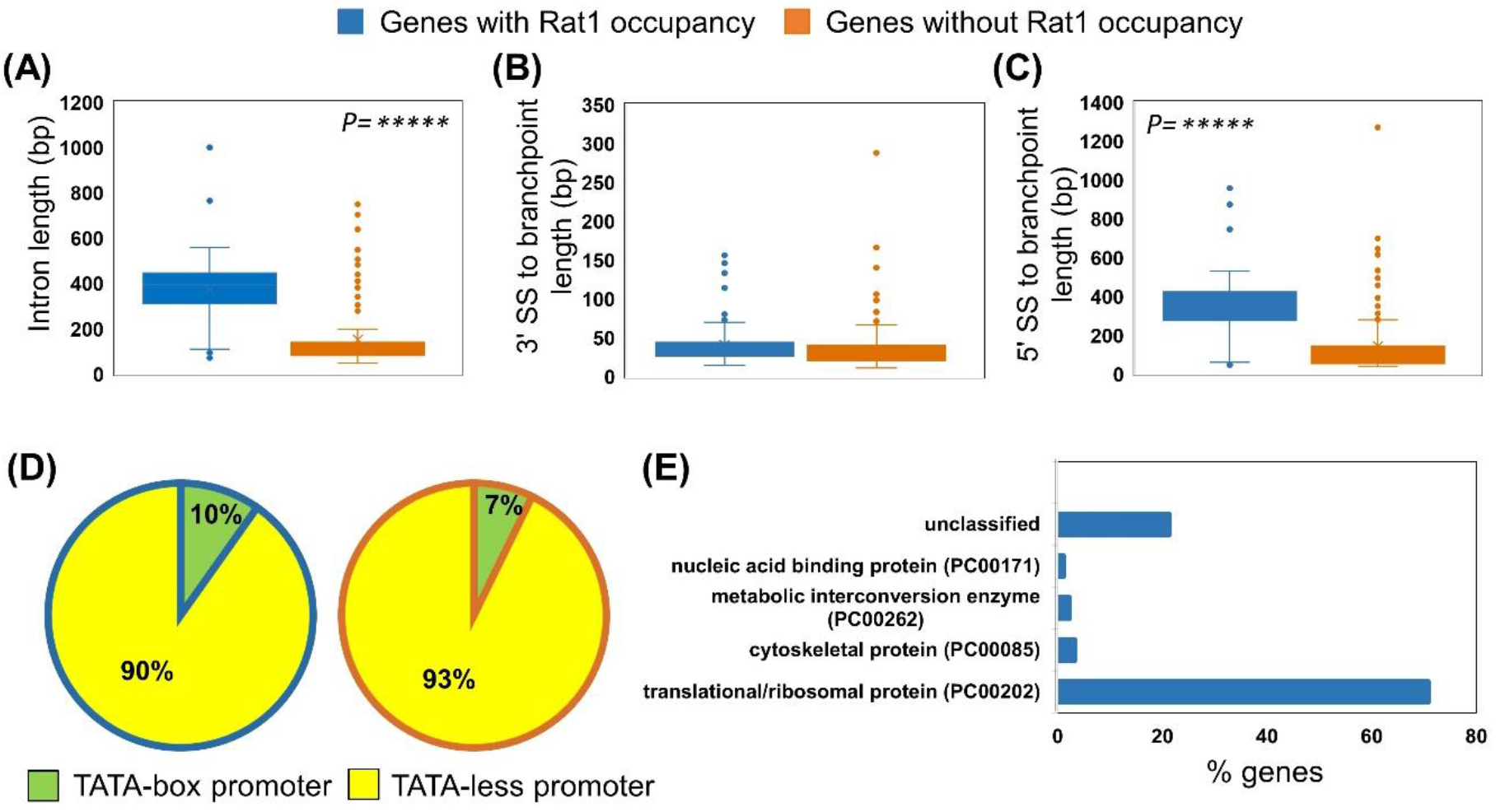
Rat1 crosslinked introns are longer in length. Genes with the presence and absence of Rat1 on intron are sorted using box-whisker plot according to intron length (A), distance between 3′ SS to branchpoint (B), distance between 5′ SS to branchpoint (C), and the promoter element TATA-box and TATA-less (D). Mann-Whitney U test was used to calculate *P value* for A, B, and C. Fisher’s exact test was used to calculate *P value* for D. Gene ontology analysis of intronic genes with Rat1 occupancy are enriched for translation/ribosomal proteins coding genes (E).

Next we searched among the 105 Rat1-occupied introns for common features, which conferred potential dependence on Rat1 for their splicing. Rat1-occupied introns do not display common predicted structures or enriched sequence motifs. Of the features that we examined, only the size of the intron showed a significant correlation (*p* value 6.9×10^−26^). On average, introns that bound Rat1 were ~400 bp long, while those not exhibiting Rat1 occupancy were ~100 bp in length (Figure 8A). The distance between branchpoint and 3′ splice site was the same for all introns (Figure 8B), but the distance between the 5′ splice site and branchpoint tended to be longer for Rat1-occupied introns (Figure 8C). Furthermore, Rat1 intron occupancy was not associated with the presence or absence of TATA-box in the promoter of the gene (*p* value 0.492) (Figure 8D). Ontological analysis revealed that the Rat1-occupied introns were enriched for ribosomal protein genes (Figure 8E).

### Rat1 interacts with splicing factors

If Rat1 is involved in splicing, it may exhibit stable or transient interactions with the splicing machinery. To determine if splicing proteins interact with Rat1, we performed biochemical purification of Rat1 employing the <u>T</u>andem <u>A</u>ffinity <u>P</u>urification (TAP) approach. Purification was performed as described in Puig et al., (2001), and the purified fraction was subjected to mass spectrometry (Figure 9–figure supplement 1). A total of 1220 proteins were identified. Rai1 and Rtt103, as expected, were present in the purified fraction of Rat1. A significant finding was the identification of five splicing proteins, Clf1, Isy1, Yju2, Sub2, and Prp43 in the Rat1 preparation (Figure 9A). Clft1, Isy1 and Yju2, however, exhibited low spectral counts. To confirm the interaction, we HA-tagged the splicing factors Clft1, Isy1, Yju2, Sub2 and Prp43 and looked for Myc-tagged Rat1 in coimmunoprecipitated fraction by Western blot. Myc-tagged Rat1 coimmunoprecipitating with all five splicing factors (Figure 9B). No signal for Rat1 was observed when the experiment was repeated in a strain without Myc-tagged Rat1 (Figure 9B). Reciprocal coimmunoprecipitation with Myc-tagged Rat1 produced similar findings but with high background. To further corroborate the interaction of Rat1 with the splicing factors described above, we repeated Rat1 coimmunoprecipitation for Prp2 and Prp16. These two yeast-specific splicing factors were not detected in our mass spectrometric analysis and no signal for Prp2 or Prp16 was detected following Rat1 pull down (data not presented). Of the five splicing factors demonstrated to interact with Rat1, Clf1, Isy1, and Yju2 are the subunits of NineTeen complex (NTC). Clf1 and Isy1 are part of the core complex and Yju2 is an accessory protein. Clf1 is an essential splicing factor that serves as a scaffold in spliceosome assembly during assembly of the tri-snRNP (U4 U5.U6). It also interacts with the branch point binding proteins Mud2 and Prp40 (Chung et al., 1999). Isy1 is not an essential splicing factor but contributes to the fidelity of the splicing reaction (Villa and Guthrie, 2005). Yju2 is an essential splicing factor that functions after Prp2 to promote the first transesterification reaction (Liu et al., 2007). Sub2, a DEAD-box RNA helicase, is a known to collaborate with Msl5 that interacts with branchpoint binding protein to promote the recruitment of U2 snRNP (Zhang and Green, 2001). Like Sub2, Prp43 is an RNA helicase; however, it functions post splicing in disassembling the U2, U5, and U6 snRNPs (Arenas and Abelson, 1997). The splicing factors, however, were not present in stoichiometric proportion to Rat1. This suggests that Rat1 is not in a stable complex with splicing factors, but transiently interacts with them during cotranscriptional splicing.

**Figure 9.**
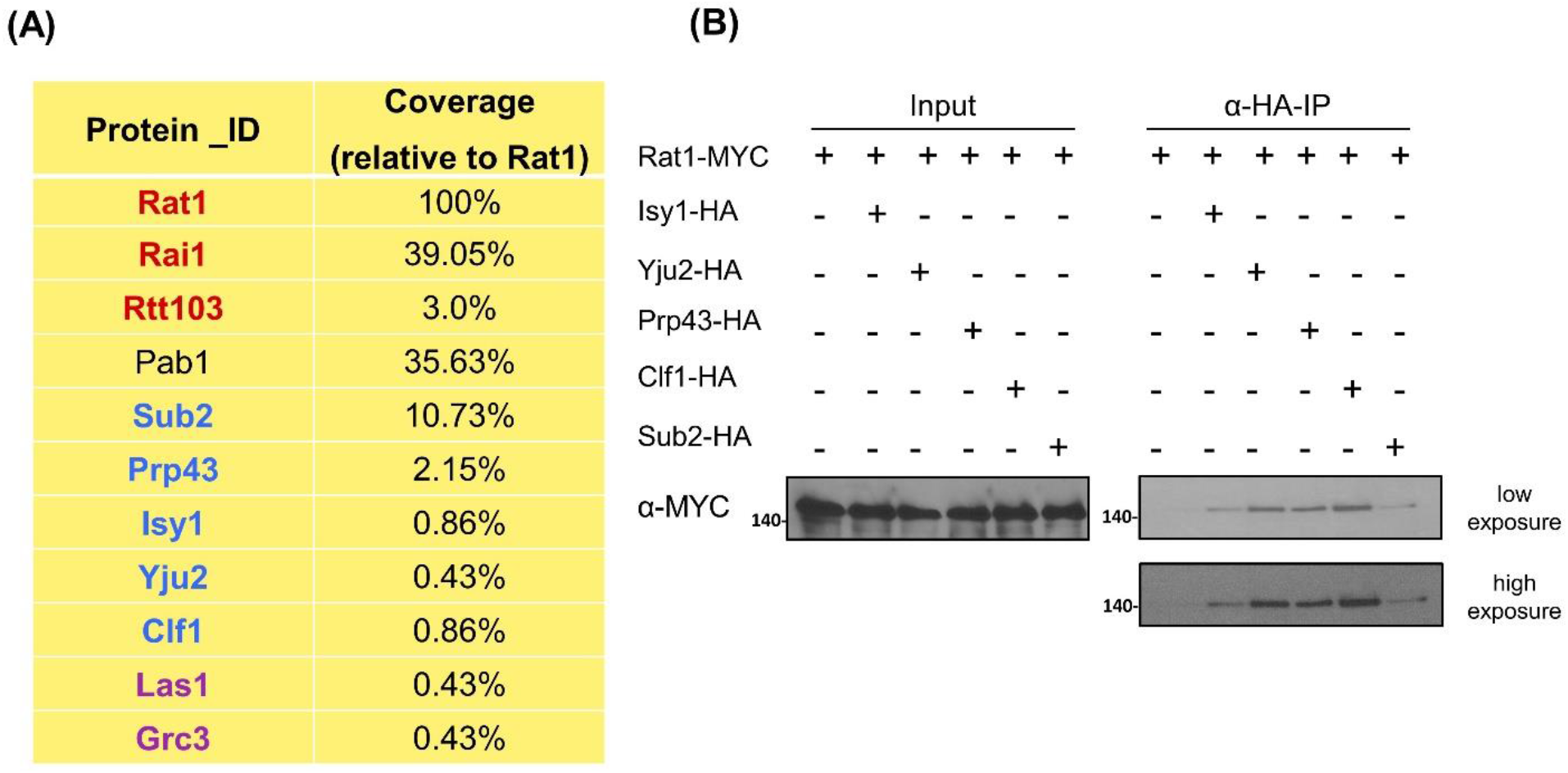
Rat1 physically interacts with splicing factors. (A) Table from mass spectrometry analysis showing presence of Rat1-interacting proteins in tandem affinity-purified Rat1 preparation. Previously known interactors of Rat1 are shown in red (Rai1 and Rtt103) and purple (Las1 and Grc3). Splicing factors (Sub2, (Prp43, Isy1, Yju2 and Clf1) are shown in blue. Rat1-interactors are listed relative to the levels of Rat1 in the preparation. (B) Coimmunoprecipitation followed by Western blot analysis confirmed Rat1 interaction with splicing factors Clf1, Isy1, Yju2, Sub2 and Prp43.

### Recruitment of splicing factor Prp2 is compromised in the absence of Rat1

Rat1 interaction with intronic sequences as well as its interaction with Isy1, Prp43, Clft1, Sub2 and Yju2 splicing factors strongly suggested a direct involvement of Rat1 in splicing of primary transcripts of a subset of genes. To further probe the role of Rat1 in splicing, we monitored the recruitment of splicing factors on introns. We reasoned that if Rat1 is indeed playing a role in splicing, recruitment of some splicing factors on the intron may be affected in the absence of Rat1. We chose Prp2 for three reasons; (1) it exhibits a genetic interaction with Rat1; (2) it is recruited on the intron before Yju2 which is the Rat1 interacting factor identified above in our analysis; and (3) its recruitment on the intron can be detected by ChIP. We checked binding of Prp2 to the intron of *APE2* by ChIP in the *Rat1-AA* strain in the presence and absence of rapamycin. Prp2-ChIP signal could be detected on the intron of *APE2, ASC1 and IMD4* in the absence of rapamycin (Figure 10, region B/C, white bars). But when Rat1 is depleted from the nucleus, the Prp2-ChIP signal registered a nearly 30-50% decline for all three genes (Figure 10, region B/C, black bars). These findings are in agreement with the observation that splicing of *APE2, ASC1 and IMD4* introns is not completely dependent on Rat1 but decreases by about 50% in the absence of Rat1 activity. Taken together, these studies strongly suggest that Rat1 has a direct role in splicing. Furthermore, Rat1 is not an essential splicing factor but enhances the efficiency of splicing of a subset of yeast genes.

**Figure 10.**
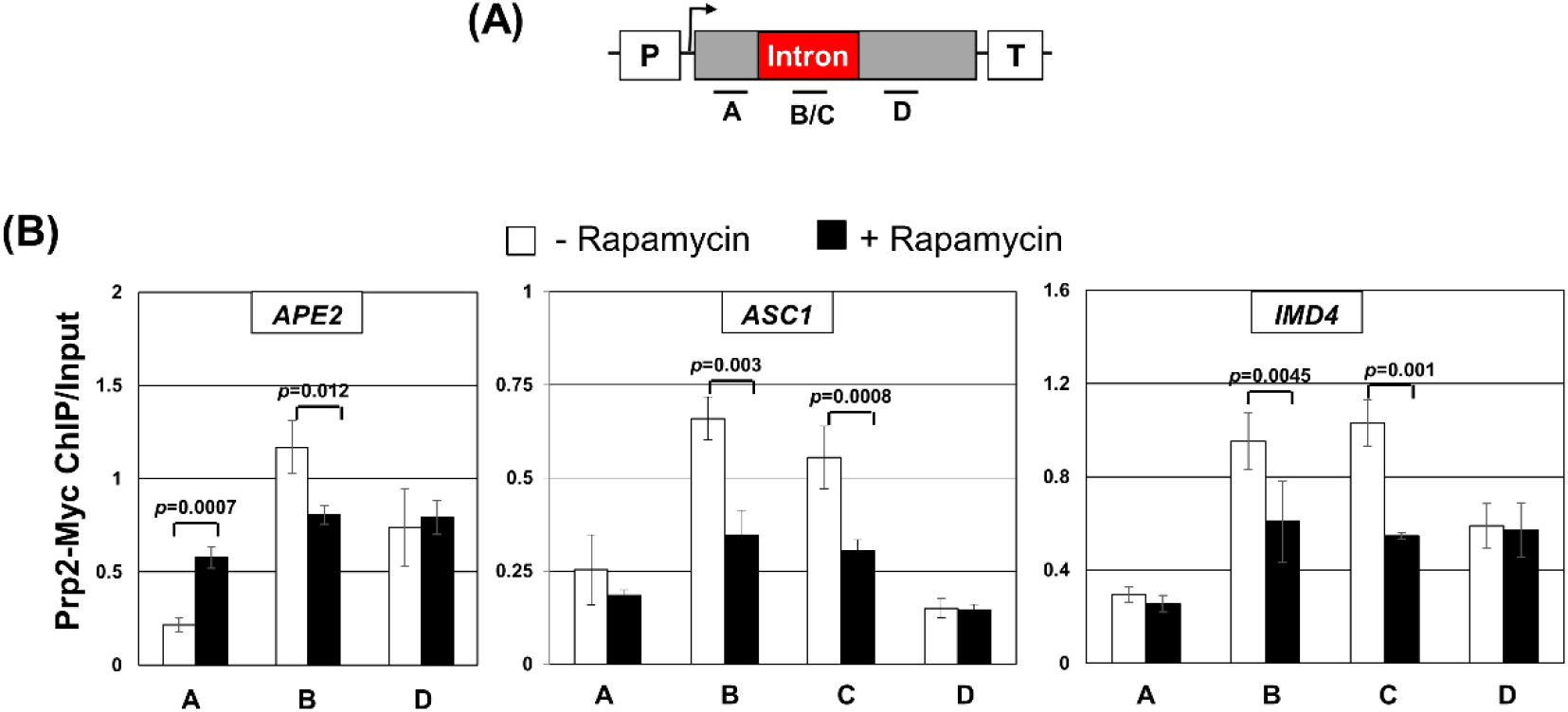
Crosslinking of Prp2 to intronic region is reduced in the absence of functional Rat1. (A) Schematic depiction of a gene indicating the position of ChIP primer pairs A, B, C and D. (B) ChIP analysis showing crosslinking of Prp2 to different regions of intron-containing genes. The input signal represents DNA prior to immunoprecipitation. The error bars represent one unit of standard deviation. The results shown are an average of two independent PCRs from two independently grown cultures.

## Discussion

Xrn2 has been implicated in degradation of unspliced mRNA in humans (Davidson et al., 2012). A report suggested involvement of Rat1 in degrading unspliced transcripts in budding yeast as well (Bousquet-Antonelli et al., 2000). In a temperature-sensitive mutant of essential splicing factor Prp2 called *prp2-1*, splicing was compromised, and unspliced transcripts could be detected at elevated temperature. In the double mutant *prp2-1/rat1-1*, there was a further increase in the amount of unspliced transcripts of selected genes. The authors attributed this additional increase in the double mutant to the role of Rat1 in degrading unspliced transcripts by its exoribonuclease activity. Estimating the level of unspliced transcripts in the *rat1-1* mutant alone, which is an important control, was not performed in this study. When we measured splicing in the *rat1-1* mutant, we observed accumulation of unspliced transcripts at the non-permissive temperature even in cells with wild type Prp2. Nuclear depletion of Rat1 by anchor-away approach gave identical results indicating that *rat1-1* mutant does not have a secondary mutation in another splicing factor. We have three pieces of evidence to support that accumulation of unspliced transcripts in the absence of Rat1 activity in yeast cells is not due to the surveillance role of Rat1 in degrading unspliced transcripts. First, unspliced transcripts were capped. Capping of unspliced mRNA was almost to the same extent as that of spliced transcripts (Figure 4). Rat1 can degrade 5′ monophosphorylated transcripts, but not 5′-7′-methylguanosine capped transcripts. Second, Rat1 does not copurify with any known decapping protein. Xrn2 in humans coimmunoprecipitates with three decapping proteins; Edc3, Dcp1a and Dcp2 (Brannan et al., 2012). Our mass spectrometric analysis of Rat1 did not yield any decapping proteins of yeast in affinity purified Rat1 preparation. Third, deletion of Rat1 interacting protein Rai1, which has been shown to remove the demethylated cap from mRNA in yeast under *in vitro* conditions (Jiao et al., 2010), has absolutely no effect on the level of unspliced transcripts in yeast cells (Figure 5–figure supplement 1). These results rule out the possibility of Rat1 degrading unspliced transcripts in budding yeast.

Rat1-dependent accumulation of unspliced mRNA can also be due to an indirect effect of Rat1 function in termination or elongation of transcription. We have four lines of evidence that unspliced transcripts are not produced because of defective termination in Rat1 mutants. First, unspliced transcripts were polyadenylated in *rat1-1* as well as *Rat1-AA* cells (Figures 1, 2 and 3). Second, in the mutant of CF1 subunit Rna14, which affects the cleavage-polyadenylation step of termination, the amount of unspliced transcripts was similar to that in isogenic wild type cells. Third, in the absence of Rat1 interacting partner Rai1, which is not a termination factor itself but affects dissociation of the polymerase from the template by stimulating 5′→3′ exoribonuclease activity of Rat1, there was absolutely no increase in the level of unspliced mRNA over the wild type control (Figure 5–figure supplement 1). Fourth, in the absence of Rtt103, which though is not an essential termination factor but has been implicated in termination (Kim et al., 2004), there was no increase in the level of unspliced mRNA over the isogenic wild type strain (Figure 5–figure supplement 1). Our results also demonstrate that generation of the unspliced transcript is not the result of increased elongation rate associated with the *rat1-1* mutant. In *Rat1-AA* strain, there was no reduction in polymerase signal over the intronic region, thereby suggesting that there was no decrease in pausing of the polymerase while transcribing the intronic region when Rat1 is nuclear depleted.

The results discussed above give credence to the hypothesis that Rat1 is playing a direct role in splicing of mRNA in budding yeast. We have three pieces of experimental evidence in support of involvement of Rat1 in splicing. First, genomewide ChIP analysis found Rat1 crosslinked to the intronic region of a number of intron-containing yeast genes. We expect a factor with a direct role in splicing to contact introns. Since splicing occurs cotranscriptionally, any protein that is involved in splicing gets indirectly crosslinked to the intronic regions on the gene through its direct contact with the transcribing RNA and the polymerase. Second, mass spectrometric analysis of tandem-affinity-purified Rat1 found at least five splicing factors, Sub2, Isy1, Prp43, Yju2 and Clf1, in the purified preparation. Three of these splicing proteins, Clf1, Isy1 and Yju2, are a part of the NineTeen complex that helps in activation of the assembled spliceosome. Third, recruitment of Prp2, an essential DEAD-box containing splicing factor in yeast, onto the intron is compromised in the absence of a functional Rat1 in the cell. Based on these results, we propose that Rat1 has a novel role in splicing of precursor mRNA in budding yeast. Rat1, however, is not an essential splicing factor as it crosslinks to less than 50% of intron containing yeast genes. Rat1 ChIP-Seq analysis revealed 105 out of 280 genes analyzed in the study showing crosslinking of Rat1 to introns. All the genes that exhibited intronic Rat1 occupancy, however, are not dependent on the protein for removal of their introns. We believe that there are other factors that determine if Rat1 localization on the intron will require it to complete the splicing reaction. In *C. elegans*, several genes that exhibited 3′ end occupancy of Rat1 did not require Rat1 for termination. Whether Rat1 facilitated termination was dictated by the promoter of the gene (Miki et al., 2017). A similar mechanism might be determining if intron bound Rat1 plays a role in splicing in yeast. It is possible that either the promoter or terminator region of the gene determines the Rat1-dependence on splicing. Even considering the genes that require Rat1 for splicing, there is no complete loss of splicing in the absence of Rat1. On average, there is 40-80% decrease in splicing efficiency in the absence of Rat1 for the genes that require it for removing an intron. These data suggest that Rat1 is not an essential splicing factor but has a rather stimulatory effect on the splicing process.

We searched for features of introns that made them dependent on Rat1 for efficient splicing. The 105 Rat1-occupying introns, which are relatively more dependent on Rat1 for splicing, were found to be approximately four-times longer in size compared to introns showing no Rat1 occupancy (Figure 8A). The longer size of Rat1-crosslinked introns was not due to a longer distance between the branchpoint and 3′ splice site, but due to a longer distance between the 5′ splice site and branchpoint (Figure 8AB and 8C). These data suggest that Rat1 probably has a role in stabilizing the interaction of the branchpoint with 5′ splice sites of long introns, which is critical for the first transesterification reaction. We propose that Rat1 can do so due to its interaction with NineTeen complex.

Apart from Prp2, Rat1 is known to exhibit a genetic interaction with at least four splicing factors; Msl5, Yhc1, Sub2 and Prp46, in yeast thereby corroborating the role of Rat1 in splicing (Bousquet-Antonelli et al., 2000; Costanzo et al., 2016; González-Aguilera et al., 2008). A logical question is if Rat1 homolog Xrn2 plays a similar role in splicing in higher eukaryotes. Xrn2 is a component of *in vivo* assembled, purified splicing complexes in human HeLa cells and chicken DT-40 cells (Chen et al., 2007). These purified supraspliceosomes contained all five U-snRNPs and other known splicing proteins. In addition, they identified some novel proteins that are not known to play any role in splicing. Xrn2 was one such protein that was present in both human and chicken supraspliceosomes. The supraspliceosomes did not contain any other termination factor besides Xrn2. The possibility of Xrn2, like Rat1, playing a role in splicing of a subset of introns in higher eukaryotes therefore cannot be ruled out.

## Materials and Methods

### Yeast Strains

The yeast strains used in this study are listed in supplemental Table S1. ZA1, ZA2, ZD47, and ZD63 were constructed by replacing the entire ORF of targeted gene by *TRP1* or *HIS3* as described in Wach, A et al., 1994. The C-terminal TAP-tagged Rat1 (ZD7) and FRB-tagged Rat1 (ZD42) strains were derived from the FY23, HHY168, and ZD42 strains by transforming with a PCR product amplified from pBS1479 and pFA6a-FRB-GFP-KanMX6, respectively. The C-terminal MYC-tagged Rat1 (ZD12) and MYC-tagged Prp2 (ZD64) strains were derived from the FY23 strain by transforming with a PCR product amplified from pFA6a-13Myc-TRP1 and pFA6a-13Myc-HIS3, respectively. The C-terminal HA-tagged Yju2 (ZD70), Clf1 (ZD71), Sub2 (ZD72), Prp43 (ZD73), and Isy1 (ZD74) strains were derived from the ZD12 (Rat1-C-MYC) strain by transforming with a PCR product amplified from pFA6a-3HA-KanMX6. The primers used for the C-terminal tagging of proteins are listed in the supplemental Table S2. A 5′ gene-specific primer and a 3′ reverse primer within the tag was used to confirm that the tag was inserted at the correct genomic position. For validation of a gene knockout, forward and reverse primer designed in the upstream and downstream regions of the open reading frame were used. Primers to confirm the strains by PCR are listed in Table S2

### Cell Culture

A 5 ml culture was started in Yeast-Peptone-Dextrose (YPD) medium using colonies from a freshly streaked plate. The culture was grown overnight at 30°C with constant shaking at 250 rpm. All *S. cerevisiae* cell cultures were grown in YPD medium unless otherwise stated. All strains except the temperature-sensitive mutants were grown at 30°C and 250 rpm. Next morning, the overnight grown cultures were diluted 1:100 in YPD broth and allowed to grow at 30°C with constant shaking until the desired A_600_ was reached. Cells were centrifuged at 1,860x g for 3 mins at 4°C and the cell pellets were used further for further experiments.

### Cell culture for Anchor-away experiment

The starter culture Rat1-FRB-tagged strain was grown in YPD broth whereas strains with plasmids expressing wild type (pRS415-WT-Rat1) or catalytically inactive mutant of Rat1 (pRS415-D235A-Rat1) were grown in synthetic complete media without leucine. The cells were incubated overnight at 30°C and 250 rpm in an orbital shaker. The cells from the starter cultures were added in 1:100 dilution to the respective culture broths. Cultures were grown until A_600_ 0.5-0.7 was reached. At this point, one-half of the culture was treated with rapamycin (Final concentration 1 mg/ml) and the other was left untreated. Both the samples were grown for another 60 min. Cells were collected by centrifugation as described above and used for RT-PCR, TRO and ChIP analysis.

### Cell culture of temperature-sensitive strains

Temperature-sensitive mutants of Rat1 (*rat1-1*) and Rna14 *(rna14-1)*, were used in this study. The overnight culture of these cells was obtained as described above except that cells were grown at 25°C. Next morning, the overnight culture was diluted 1:50, and not 1:100 as is normally done, in YPD broth. The cells were grown with gentle shaking at permissive temperature (25°C) until the A_600_ reached 0.25 and 0.4 for *rat1-1* and *rna14-1* strains respectively. Cells were then transferred to non-permissive temperature (37°C) for 3 hours (*rat1-1)* or 1 hr (*rna14-1*). Cells were harvested at A_600_ between 0.5-0.7 by centrifugation at 3000 rpm for 3 mins at 4°C. Another cell culture of these strains was run in parallel, incubated at permissive temperature (25°C) until desired A_600_ 0.5-0.7 reached. An isogenic wild type strain was subjected to similar growth conditions and the final cell density was normalized considering the mutants strain cell density. The harvested cells were used for RT-PCR, TRO and RNA-IP analysis and were processed as described below in the respective experimental sections.

### Isolation and analysis of RNA

Cells were grown and processed as discussed in previous section. RNA was extracted as described in(McNeil and Smith, 1986). 2 μg/μl of RNA was used for cDNA synthesis using M-MuLV reverse transcriptase with oligo-dT and 5S 3′ primers, respectively. cDNA was diluted ten times prior to PCR amplification by Taq DNA polymerase. The gene specific primers used for PCR reaction are listed in Table S2. Each PCR signal was normalized with respect to 5S ribosomal RNA control. In parallel, a negative control without reverse transcriptase was run to ensure DNA contamination is not contributing to any RT-PCR signal. Each experiment was repeated with at least three independently grown cultures.

### Transcription Run-On Assay (TRO)

FRB-tagged strains were processed as discussed in previous section. The transcription run-on assay was performed as described in (Dhoondia et al., 2017). For the transcription analysis, 500 ng/μl of RNA was used for cDNA synthesis with Superscript IV reverse transcriptase using oligo-dT and 5S 3′ primers. cDNA was diluted five times prior to PCR amplification by Platinum SuperFi DNA polymerase. The gene specific primers used for PCR reaction are listed in Table S2. Each PCR was normalized with respect to 5S ribosomal RNA control. A sample without reverse transcriptase enzyme was run in parallel as a negative control. Each experiment was repeated with at least three independently grown cultures.

### Chromatin Immunoprecipitation (ChIP)

100 ml of cell culture was treated with 1% formaldehyde and 20 mM disuccinimidyl glutarate (DSG) (20593; ThermoFisher Scientific) in 1X PBS with constant shaking for 20 minutes at 25°C. Crosslinking was quenched by the addition of 125 mM glycine and incubating for 5 minutes at 25°C. Cells were lysed and chromatin was isolated and sonicated as described previously (El Kaderi et al., 2009). ChIP for Rat1-TAP was performed by incubating 500 μl of the solubilized chromatin sample with 20 μl of IgG Sepharose 6 Fast Flow (GE healthcare) for 3 hours at 4°C. For Prp2-Myc ChIP, 400 μl of the solubilized chromatin was pre-incubated with 5 μl Anti-Myc antibody (PA1-981; Invitrogen) for 2 hours at 4°C. 20 μl of Magnetic Dynabeads Protein G was prewashed two times prior to use. Antibody-antigen complex was captured on magnetic Dynabeads (1003D) by incubating at room temperature for 15 minutes. The tubes were placed on a ThermoFisher Scientific Magjet rack, and supernatant containing unbound antigen and antibody was discarded. After removing the unbound material, the Sepharose or magnetic beads were subjected to a series of washing steps and processed to obtain DNA as described in El Kaderi et al., 2009. Purified DNA from the input and ChIP fraction was used as a template for PCR using the primers specific for a gene. The primers used for ChIP-PCR analysis are listed in Table S2. The association of a protein to a given genomic region was expressed as ChIP over input ratios and subjected to quantification and statistical analysis as described in (El Kaderi et al., 2012). Each experiment was repeated with at least three independently grown cultures.

### Data mining and re-analysis of published datasets

Published ChIP-Seq analysis dataset from the study by Baejen et al. (2017) was retrieved through the Gene Expression Omnibus (GEO), GSE79222. Raw reads were further processed and analyzed using the Galaxy web platform at usegalaxy.org (Afgan et al., 2016). Raw reads (50 bp paired end reads) were aligned to the *S. cerevisiae* genome (sacCer3, version 64.2.1) using the Bowtie (version 2.3.4.2) (Langmead and Salzberg, 2012) following the parameters described in Baejen et al., 2017. To normalize and compare two BAM files and obtain a log2 ratio of IP/Input in the form of a bigwig file, bamCompare (Galaxy Version 3.1.2.0.0) in deepTools (Ramírez et al., 2016) was used. The aligned reads with MAPQ alignment quality smaller than 7 (-q 7) were skipped. Signal extraction scaling (SES) factor was used for scaling and establishing ChIP enrichment profile over log2 scale with the options: –l 100 –n 100000 and the bin size of 20. The subsequent data in the bigwig file was selectively enriched for the genomic regions in the BED file using computeMatrix function (Galaxy Version 3.3.0.0.0). Specifically, three BED files were used in this study; 280 intron-containing genes with TSS and TES coordinates, 280 intron-containing genes with intron start and intron end coordinates, and 2392 non-intronic genes with TSS and TES coordinates. Heatmap was plotted to visualize the score distributions of the transcription factors or RNAPII enrichment associated with genomic regions specified in the BED files using the plotHeatmap (Galaxy Version 3.3.0.0.1). The metagene plot of the transcription factors and RNAPII was drawn with the plotProfile function (Galaxy Version 3.3.0.0.0).

ChIP/Input data in bigwig files were imported to Integrated Genome Browser (IGB) (Nicol et al., 2009) for visualization. In case of RNAP II heatmap and metagene profile, the occupancy pattern was plotted on the window of intron scaled to 400 bp and 0.4 kbp upstream and 1.0 kbp downstream of the intronic region. For transcription factors, Rat1 and Pcf11, the plot was constructed on the window of introns scaled to 400 bp and 0.1 kbp upstream and 1 kbp downstream of the intronic region. A confidence interval of mean is computed at each bins (20 bp bin sizes) using bootstrap sampling or Rat1 and Pcf11 and is visually represented by a semi-transparent color with a width corresponding to one unit of the standard error. The correlation of Rat1 intron occupancy for the length of the intron, distance from 5′ splice site to branch point, and distance from 3′ splice site to branch point has been determined for 280 intronic genes using box-whisker plots. Mann-Whitney U test was used to determine statistical significance of Rat1 occupancy for the intron length, distance from 5′ or 3′ splice site to the branch point. The list of intron-containing genes with TATA box or TATA-less promoters were filtered from data file of yeast genes characterized into TATA-containing or TATA-less promoter (Basehoar et al., 2004). Distribution of TATA box or TATA-less promoter reflecting the percentage of intron containing genes with and without enrichment of Rat1 on the intronic region is shown using a pie chart. The statistical significance for the promoter preference and Rat1 occupancy was determined using Fisher’s exact test. Statistical analysis was computed on Graphpad Prism 8.4.2. Gene Ontology (GO) analyses using SGID of the intronic genes with Rat1 occupancy was performed using PANTHER (http://geneontology.org/) using default options (Ashburner et al., 2000). Additionally, we performed GO ontology functional analysis and detection of protein class of the genes in the dataset. Benjamini-corrected *P-value* was used for ranking GO terms.

### RNA Immunoprecipitation

Mutant *rat1-1* strain and the isogenic wild type strain were grown, and RNA was isolated as described previously. 2 μg/μl of RNA was set aside as input that was used to quantify the total amount of spliced and unspliced RNA in the samples. Briefly, cDNA synthesis of 2 μg/μl of RNA was performed using Superscript IV reverse transcriptase and oligo-dT 3′ primer. cDNA was diluted five times prior to use in PCR amplification using Platinum SuperFi DNA polymerase. The gene specific primers used for PCR reaction are listed in Table S2.

The immunoprecipitation of the capped transcripts was performed by incubating 4 μl of Anti-7′-methylguanosine (m^7^G)-Cap mAb (RN016M; MBL) with 40 μg of total RNA adjusted to the total volume of 400 μl using RNA-IP binding buffer (20 mM HEPES KOH, 10 mM MgCl_2_, 150 mM NaCl, 0.5 mM DTT, 80U RNase inhibitor). 15 μl of prewashed magnetic Dynabeads was used to capture immunoprecipitation complex by incubating for 15 mins at room temperature. Unbound material was discarded. The beads were washed three times with 400 μl RNA-IP binding buffer. After the final wash, 100 μl of binding buffer was added, mixed to resuspend the beads, and transferred to a new tube. The buffer was removed as described above. Capped RNA bound to the Anti-7′-methylguanosine (m^7^G)-Cap mAb was released by 100 μl of elution buffer (20 mM HEPES-KOH, 1% SDS, 2mM EDTA). RNA elution was performed by gently shaking tubes on a nutator for 5 minutes at room temperature. Beads were separated using magnetic rack and the supernatant containing capped RNA was transferred into a new tube and subjected to phenol-chloroform extraction. RNA was precipitated using 0.1 volume of 3 M sodium acetate, 2.5 volume of 100 % ethanol and glycogen as a carrier. The RNA pellet was collected by centrifugation at 16,168 x g for 10 minutes at 4 °C, air-died and resuspended in 16 μl of RNase-free water. The concentration of RNA was determined using Nanodrop. Approximately 200 ng/μl of RNA was used for cDNA synthesis by M-MuLV reverse transcriptase (NEB) and 3′ oligo-dT primer. cDNA was diluted two times prior to PCR amplification. The gene specific PCR primers used are listed in Table S2. A negative control without reverse transcriptase enzyme was carried out in parallel to rule out the possibility of DNA contamination.

### Preparation of protein extract from yeast cells

The Rat1-TAP and an isotype no-tag control strains were cultured in YPD media to an A_600_ of ~4.0. Two liters of each cultures were harvested by centrifugation at 5000 rpm for 5 minutes at 4°C. Cell pellets were washed with 1xTBS and resuspended in 15 ml of lysis buffer (20 mM HEPES-KOH, 10 mM MgCl_2_, 150 mM NaCl, 1 mM PMSF, 0.5 mM DTT and 10% glycerol). The resultant cell suspension was added dropwise to liquid nitrogen filled 50 ml conical tube and stored at −80 °C.

Lysis of the frozen cell beads was performed in Retsch MM301 mixer mill. The mixer mill chamber and stainless-steel ball were chilled in liquid nitrogen before use. The frozen cell beads and stainless-steel ball were transferred to the chamber, secured properly and placed on the holder of mixer mill. The mixer mill was set for 3 mins cycles at 15 Hz. Lysis was performed by repeating this cycle for 15 times. After each cycle, the chambers were unfastened from the holder and submerged into liquid nitrogen to ensure that frozen cells are not thawing. After completion of 15 lysis cycles, lysed cells were scrapped from the chambers and transferred to a 50 ml conical tube. An additional 30 ml of lysis buffer was added to obtain a paste of lysed yeast cells. This suspension was centrifuged at 5000 rpm for 10 mins at 4°C to remove cell debris. The supernatant represents the whole cell extract. Soluble extract was obtained from the whole cell extract as described in Svejstrup et al., (2003) and stored at −80°C until further use.

### Purification of Rat1 complex

The Rat1 complex was purified as described in Puig et al., 2001 with slight modifications. Immunoprecipitation was carried out in the Poly-Prep® Chromatography columns (BIORAD). In the first affinity purification step, 600 μl of IgG Sepharose® 6 Fast Flow (GE healthcare) bead suspension was taken into a column and washed once with 10 ml of IP binding buffer (20 mM HEPES-KOH pH 7.9, 10 mM MgCl2, 150 mM NaCl, 10% Glycerol, 0.5 mM DTT, 1 mM PMSF). Three such columns were run for each sample. The soluble extract obtained from yeast cells was added to the column and incubated with the IgG-Sepharose beads on a nutator for 3 hours at 4°C. Washing and elution steps were performed by gravity flow. The unbound sample was drained and a 100 μl aliquot of this sample is set aside as flow through. The beads are washed thrice with 10 ml wash buffer to ensure that any unbound proteins are not carried forward in the analysis. A wash with 10 ml of TEV cleavage buffer (20 mM Tris-HCl pH 8.0, 150 mM NaCl) was followed to equilibrate beads with the buffer for TEV cleavage step. Cleavage at TEV site in TAP tag was carried out using 160 units of TEV protease (MC Labs) in 1 ml of TEV cleavage buffer. The column was incubated at 16°C for 2 hours. The eluate from this enzymatic elution step is collected in a new tube. An additional 500 μl of TEV cleavage buffer was added to collect 1.5 ml of the total eluate from this step.

About 300 μl of Calmodulin-Sepharose 4B (GE healthcare) beads suspension was equilibrated with 10 ml of calmodulin-binding buffer (50 mM Tris-HCl pH 8.0, 150 mM NaCl, 1 mM magnesium acetate, 1 mM imidazole, 2 mM β-mercaptoethanol and 3 mM CaCl_2_). To the previously collected TEV eluate, 4.5 ml of calmodulin-binding buffer and 4 mM CaCl_2_ were added. This sample is then transferred to a column that contains equilibrated Calmodulin-Sepharose 4B beads. The binding step was carried out for 2 hours at 4°C. Unbound proteins were drained from the column under gravity and 500 μl was set aside as flow through for further analysis. Protein captured on Calmodulin-Sepharose 4B were washed thrice with 10 ml of calmodulin-binding buffer. The bound proteins and their associated partners were eluted in five fractions of 200 μl each using calmodulin-elution buffer (50 mM Tris-HCl pH 8.0, 150 mM NaCl, 1 mM magnesium acetate, 1 mM imidazole, 2 mM β-mercaptoethanol and 3 mM EGTA). Trichloroacetic acid (TCA) was added at a final concentration of 30% to the eluate from the second affinity purification steps. The samples were incubated at 4°C for 2 hours on ice. Samples were then centrifuged at 15,700x g for 30 minutes at 4°C to pellet precipitated proteins. The supernatant was carefully removed, and any traces of TCA were removed by spinning again at 15,700x g for 1 minute at 4°C. The TCA precipitated protein pellet was washed with 500 μl of 100% acetone. The acetone washed pellet was air-dried at room temperature for 10 minutes and then resuspended in 40 μl of buffer (100 mM Tris-HCl pH 8.8, 1% SDS).The presence of Rat1-TAP at different steps of purification was analyzed using western blot analysis as described in El Kaderi et al., 2009.

### Silver Staining

Silver staining was performed to visualize protein bands present in the eluate fraction after the second affinity purification step of the Tandem-Affinity-Purification approach (TAP). The enrichment of the bait protein, Rat1, and its associated interacting partners was monitored after each step. The electrophoresed gel was fixed in 150 ml solution of 50% methanol and 5% acetic acid by gently shaking the gel at room temperature for 20 min. The gel was then washed with 150 ml 50% methanol, followed by washing with water for 10 mins. Both steps were carried out on a shaker at room temperature. The gel was sensitized in a solution containing 0.02% sodium thiosulfate for 1 min at room temperature. The gel was rinsed for 1 min with 150 ml of water and incubated in 0.1% silver nitrate and 0.08% formaldehyde for 20 mins on a shaker. The gel was further rinsed twice with 150 ml of water to remove unbound silver nitrate. The protein bands were visualized using a developer solution that comprised of 2% sodium carbonate and 0.04% formalin (37%). The developer was replaced once solution turned yellow. The bands were allowed to develop until the desired intensity was obtained. The developing reaction was stopped by washing the gel for 10 min with 150 ml 5% acetic acid. The gel was washed with water once before recording the image on ChemiDoc MP (*BIO-RAD*).

### Mass Spectrometry analysis

Three fractions of the proteins eluted from the Calmodulin-Sepharose column were pooled together and submitted to the proteomic core facility at Wayne State University. The processing of samples from the wild type Rat1-TAP (test) and wild type no-tag (control) samples was performed at the proteomic facility as described below. To gauge the background contamination due to sample processing, a background (bkg) tube without protein was processed in parallel with the sample tubes. Samples were precipitated with two volumes of ice-cold 100% methanol in 1 mM acetic acid at −20°C overnight. The next day, samples were spun at 17,000x g for 20 min at 4°C and the supernatant was removed. The resultant pellets were rinsed with 100 μl methanol-acetic acid. Samples were dried in Speed-Vac for 3 min. The pellets were solubilized in 50 mM triethylammonium bicarbonate (TEAB) using Qsonica sonicator, then reduced with 5 mM DTT and alkylated with 15 mM iodoacetamide (IAA) under standard conditions. Excess IAA was quenched with an additional 5 mM DTT. The samples were then digested overnight in 50 mM TEAB with sequencing-grade trypsin (Promega). The next day, digests were acidified with 1% formic acid and a 10% aliquot of the supernatant was analyzed.

The peptides were separated by reversed-phase chromatography (Acclaim PepMap100 C18 column, Thermo Scientific), followed by ionization with the Nanospray Flex Ion Source (Thermo Scientific), and introduced into a Q Exactive™ mass spectrometer (Thermo Scientific). Abundant species were fragmented with high-energy collision-induced dissociation (HCID). Data analysis was performed using Proteome Discoverer 2.1 (Thermo Scientific) which incorporated the SEQUEST algorithm (Thermo Scientific). The Uniprot_Yeast_Compl_20160407 database was searched for yeast protein sequences and a reverse decoy protein database was run simultaneously for false discovery rate (FDR) determination. The data files were loaded into Scaffold (Proteome Software) for distribution. SEQUEST was searched with a fragment ion mass tolerance of 0.02 Da and a parent ion tolerance of 10 PPM. Carbamidomethylation of cysteine was specified in SEQUEST as a fixed modification. Deamidation of asparagine and glutamine, oxidation of methionine, and acetylation of the N-terminus were specified in SEQUEST as variable modifications. Minimum protein identification probability was set at <= 1.0% FDR with 2 unique peptides at <= 1.0% FDR minimum peptide identification probability. (0.5% protein decoy FDR, 0.16% peptide decoy FDR).

### Co-immunoprecipitation

Cells expressing Rat1-C-MYC and Clf1/Isy1/Prp43/Sub2/Yju2-C-HA were grown in 100 ml of YPD to an OD_600_ of 1.2-1.4, harvested, washed with 1x Tris-buffer Saline, and then suspended in 1.0 ml of lysis buffer (20 mM HEPES-KOH, 10 mM MgCl_2_, 150 mM NaCl, 1 mM PMSF, 0.5 mM DTT and 10% glycerol). The cell suspension was flash frozen in liquid nitrogen and processed as described in El Kaderi et al 2009. For immunoprecipitation, the cell lysate was incubated with 20 μl of Anti-HA Agarose (Pierce). Beads were washed thrice with lysis buffer and elution was carried out in SDS sample buffer lacking 2-mercaptoethanol. Western blot analysis was performed using the anti-MYC (PA1-981; Invitrogen) to detect the presence of Rat1-C-MYC.

## Acknowledgments

This work was supported by grant from National Science Foundation (MCB1936030) to AA. The content is solely the responsibility of the authors and does not necessarily represent the official views of the National Science Foundation. We thank Dr. Claire Moore of Tufts University for kindly providing the plasmids expressing wild type and mutant Rat1.We are grateful to Dr. Victoria Meller and Dr. Jared Schrader of Wayne State University for critical reading of the manuscript. We thank lab members for helpful suggestions.

## Competing interests

The authors declare that they have no known competing financial interests or personal relationships that could have appeared to influence the work reported in this paper.

---

**Figure 4–figure supplement 1.**
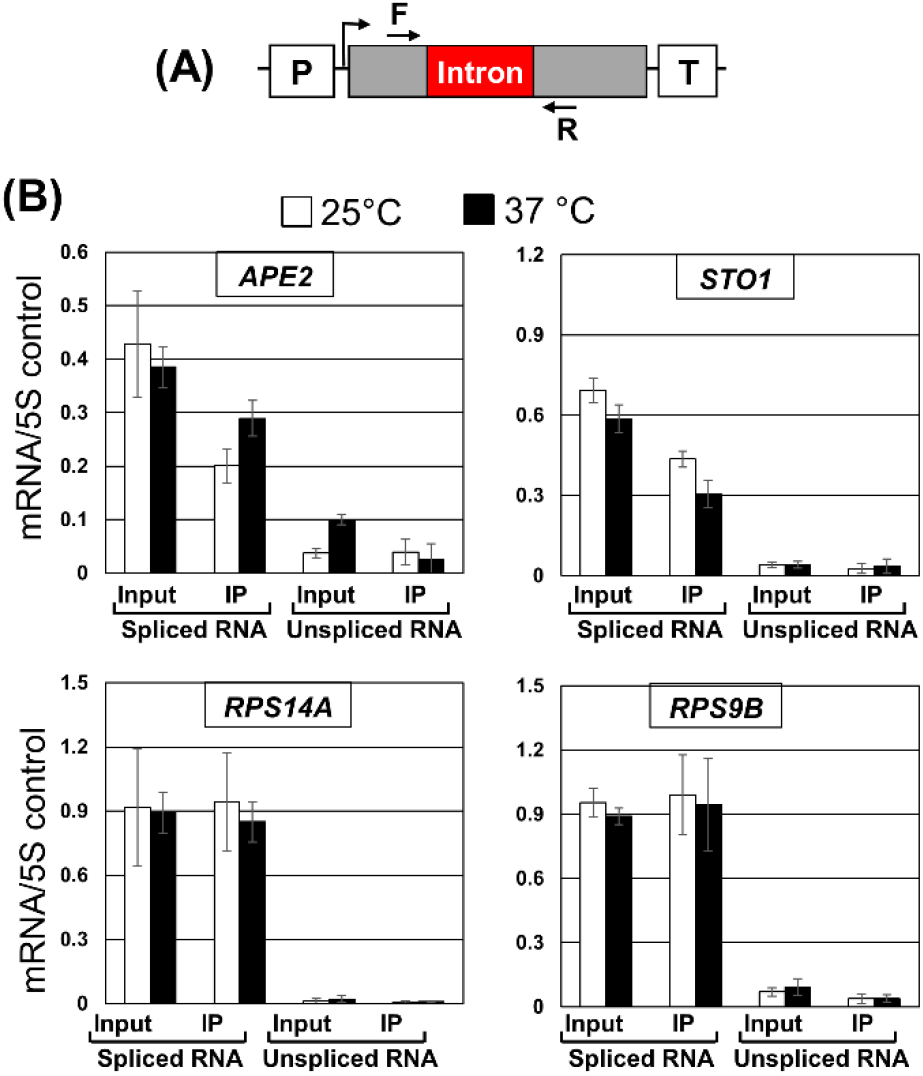
RNA immunoprecipitation analysis shows that spliced and unspliced transcripts in wild type strain (FY23). (A) Schematic depiction of a gene showing the position of primers F and R used in RT-PCR analysis. (B) Affinity purified RNA using anti-m7G were reverse transcribed. Quantification of data shown in for the indicated genes in wild type (WT) at the indicated temperatures. 5S rRNA was used as normalization control.

**Figure 5–figure supplement 1.**
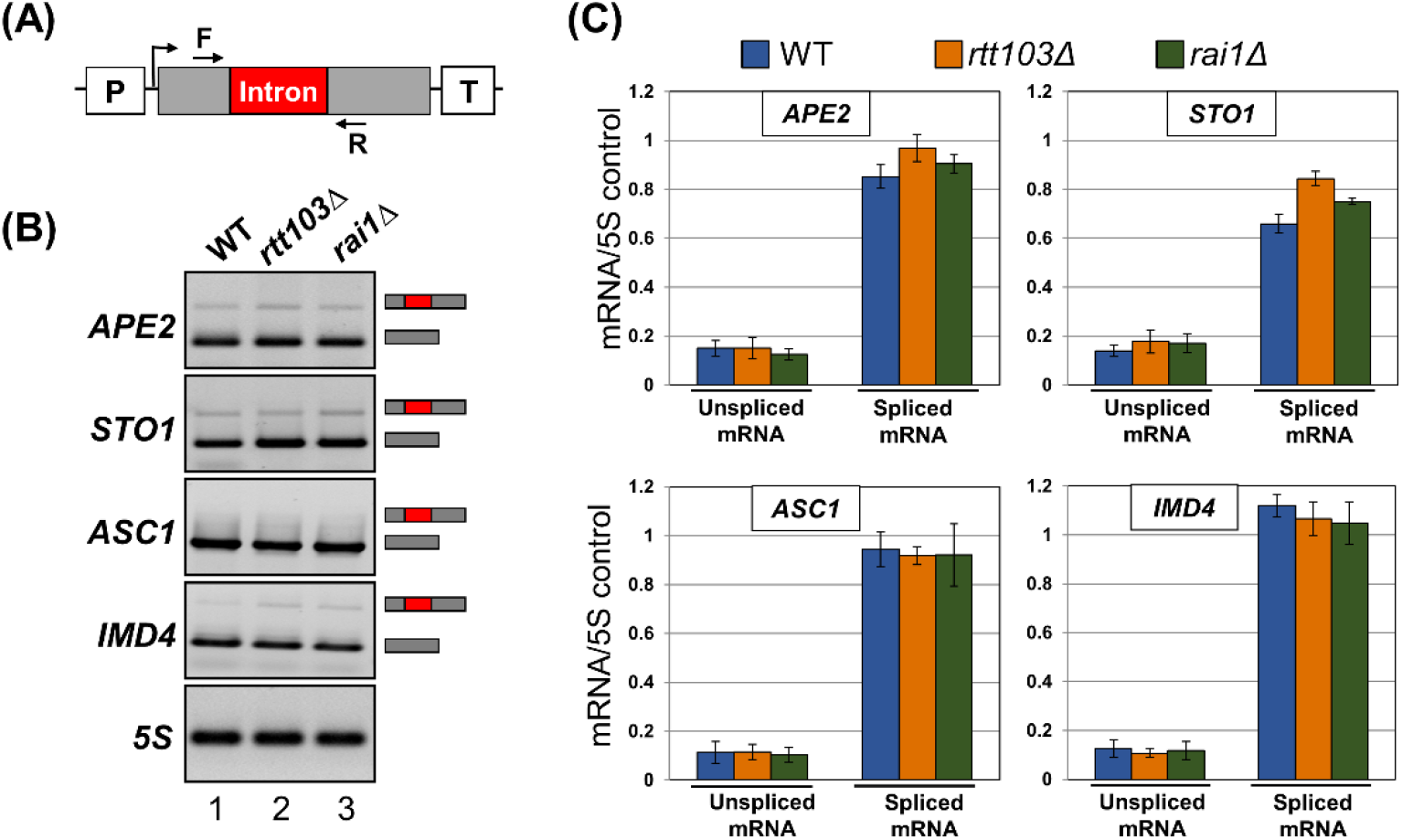
Rat1 termination complex is not required for accumulation of unspliced transcripts. (A) Schematic depiction of a gene showing the position of primers F and R used in RT-PCR analysis. (B) Gel pictures showing RT-PCR products for the indicated genes in wild type (WT), *rtt103Δ* and *rai1Δ* strains. (C) Quantification of data shown in (B). 5S rRNA was used as normalization control.

**Figure 7–figure supplement 1.**
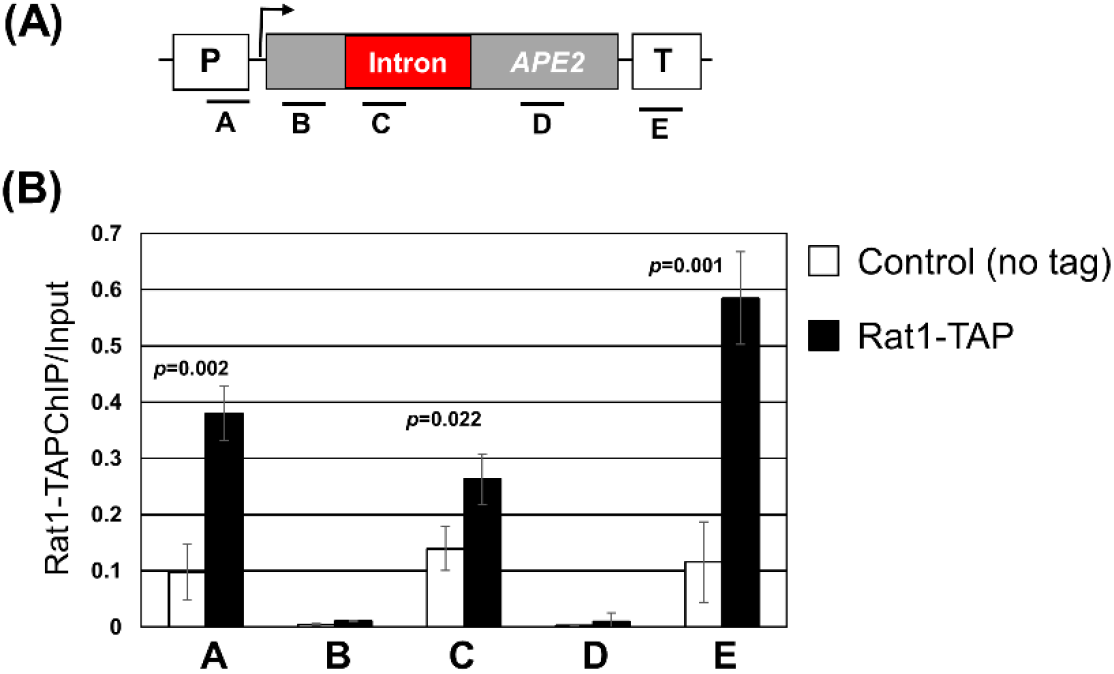
Crosslinking of Rat1 to intron of *APE2* gene. (A) Schematic depiction of *APE2* indicating the position of ChIP primer pairs A, B, C, D, and E. (B) Quantitative analysis showing crosslinking of Rat1 to different regions of *APE2* in Rat1-TAP and WT (no tag) strains represented using black and white bars, respectively. The input signal represents DNA prior to immunoprecipitation.

**Figure 7–figure supplement 2.**
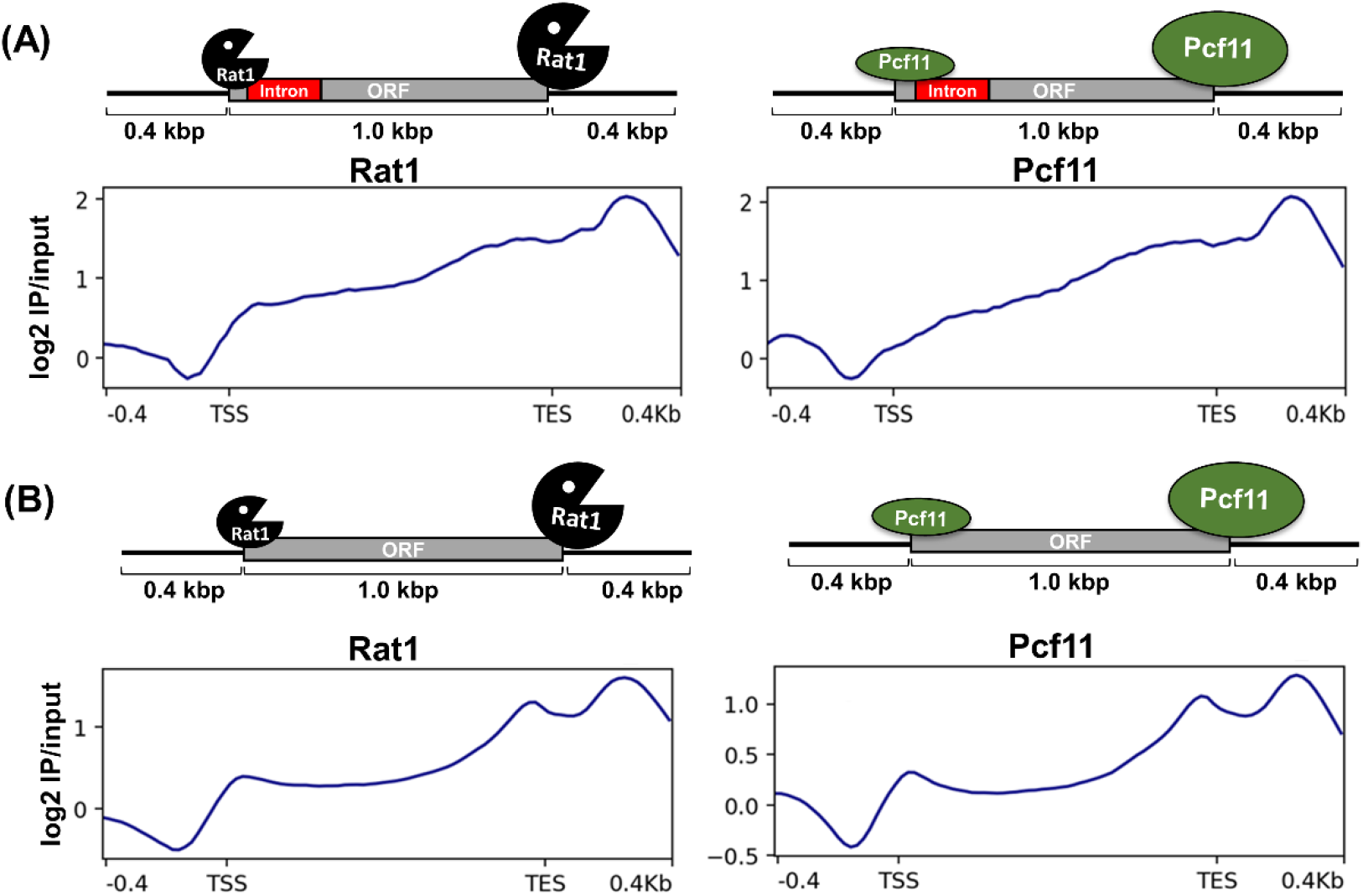
Rat1 occupancy profile differs from Pcf11 at the intronic genes and not at the non-intronic genes. Average normalized plot of Rat1 and Pcf11 occupancy in WT strain, mapped to 2392 genes that are non-intronic (A) and 280 genes that are intronic. The metagene plots are are constructed with the coordinates of transcription start site (TSS) and transcription end site (TES) and distance of the open reading frame (ORF) averaged to 1 Kbp.

**Figure 8–figure supplement 1.**
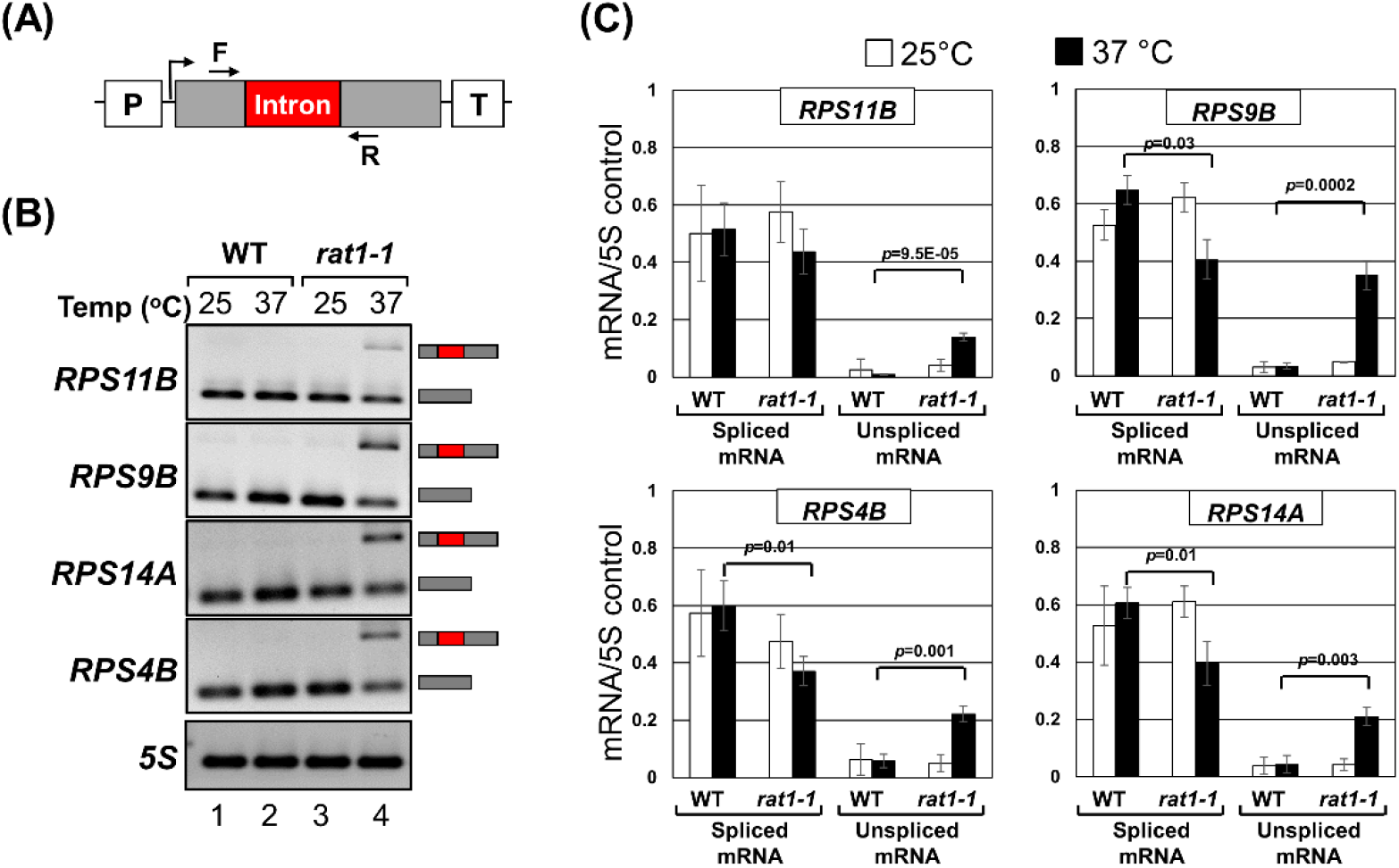
Rat1 intronic occupancy correlates with Rat1-dependent accumulation of unspliced transcripts in *rat1-1* mutant at non-permissive temperature. (A) Schematic depiction of a gene showing the position of primers F and R used in RT-PCR analysis. (B) Gel pictures showing RT-PCR products for the indicated genes in wild type (WT) and *rat1-1* mutant cells at the indicated temperatures. (C) Quantification of data shown in (B). 5S rRNA was used as normalization control.

**Figure 9–figure supplement 1.**
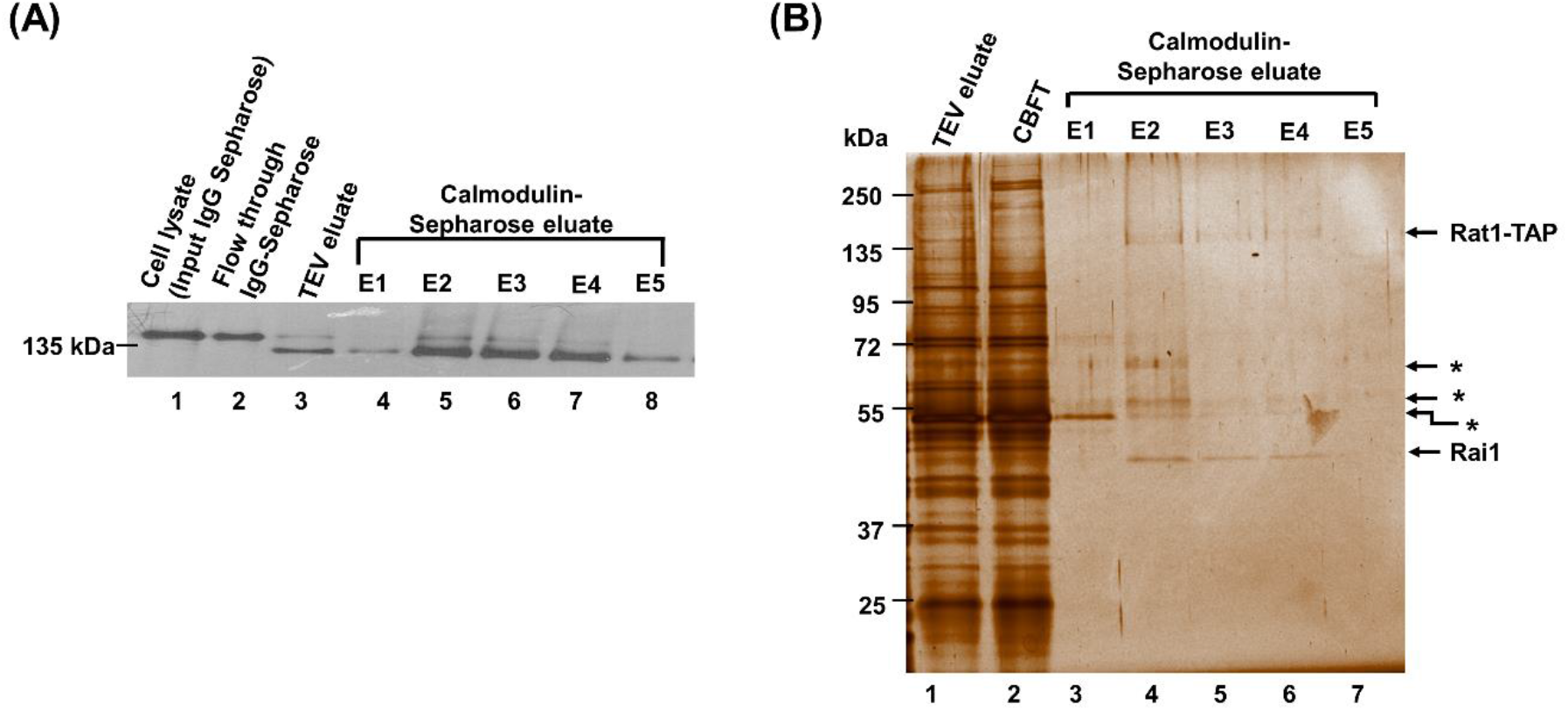
Purification of proteins interactors of Rat1 from yeast strain with TAP tag inserted at the C-terminus of Rat1 gene. (A) Western blot analysis at the different steps of the purification tracks Rat1 before and after TEV cleavage step. Rat1 is higher molecular weight prior TEV cleavage (lane 1 and 2) compare to post TEV cleavage (lane 3-8). (B) Silver stained gel (10% SDS-PAGE) showing the presence of proteins in after TEV digestion (TEV eluate), flow through from calmodulin binding step (CBFT), and five elution fractions (E1-E5) following calmodulin binding step. High amount of proteins in TEV and CBFT represents considerable contaminant proteins. Eluate fractions E2, E3, E4 represents elution Rat1 and associated proteins in a complex.

## Notes

### Competing Interest Statement

The authors have declared no competing interest.

